# *BARD1* germline variants induce haploinsufficiency and DNA repair defects in neuroblastoma

**DOI:** 10.1101/2023.01.31.525066

**Authors:** Michael P. Randall, Laura E. Egolf, Zalman Vaksman, Minu Samanta, Matthew Tsang, David Groff, J. Perry Evans, Jo Lynne Rokita, Mehdi Layeghifard, Adam Shlien, John M. Maris, Sharon J. Diskin, Kristopher R. Bosse

## Abstract

**Importance:** High-risk neuroblastoma is a complex genetic disease that is lethal in 50% of patients despite intense multimodal therapy. Our genome-wide association study (GWAS) identified single-nucleotide polymorphisms (SNPs) within the *BARD1* gene showing the most significant enrichment in neuroblastoma patients, and also discovered pathogenic (P) or likely pathogenic (LP) rare germline loss-of-function variants in this gene. The functional implications of these findings remain poorly understood.

**Objective:** To define the functional relevance of *BARD1* germline variation in children with neuroblastoma.

**Design:** We correlated *BARD1* genotype with *BARD1* expression in normal and tumor cells and the cellular burden of DNA damage in tumors. To validate the functional consequences of rare germline P-LP *BARD1* variants, we generated isogenic cellular models harboring heterozygous *BARD1* loss-of-function (LOF) variants and conducted multiple complementary assays to measure the efficiency of DNA repair.

**Setting:** (N/A)

**Participants:** (N/A)

**Interventions/Exposures:** (N/A)

**Main Outcomes and Measures:** *BARD1* expression, efficiency of DNA repair, and genome-wide burden of DNA damage in neuroblastoma tumors and cellular models harboring disease-associated *BARD1* germline variants.

**Results:** Both common and rare neuroblastoma associated *BARD1* germline variants were significantly associated with lower levels of *BARD1* mRNA and an increased burden of DNA damage. Using neuroblastoma cellular models engineered to harbor disease-associated heterozygous *BARD1* LOF variants, we functionally validated this association with inefficient DNA repair. These *BARD1* LOF variant isogenic models exhibited reduced efficiency in repairing Cas9-induced DNA damage, ineffective RAD51 focus formation at DNA doublestrand break sites, and enhanced sensitivity to cisplatin and poly-ADP ribose polymerase (PARP) inhibition.

**Conclusions and Relevance:** Considering that at least 1 in 10 children diagnosed with cancer carry a predicted pathogenic mutation in a cancer predisposition gene, it is critically important to understand their functional relevance. Here, we demonstrate that germline *BARD1* variants disrupt DNA repair fidelity. This is a fundamental molecular mechanism contributing to neuroblastoma initiation that may have important therapeutic implications, and these findings may also extend to other cancers harboring germline variants in genes essential for DNA damage repair.

**Key Points:** *Question:* How do neuroblastoma patient BRCA1-associated RING domain 1 (*BARD1*) germline variants impact DNA repair?

*Findings:* Neuroblastoma-associated germline *BARD1* variants disrupt DNA repair fidelity. Common risk variants correlate with decreased *BARD1* expression and increased DNA double-strand breaks in neuroblastoma tumors and rare heterozygous loss-of-function variants induce *BARD1* haploinsufficiency, resulting in defective DNA repair and genomic instability in neuroblastoma cellular models.

*Meaning:* Germline variation in *BARD1* contributes to neuroblastoma pathogenesis via dysregulation of critical cellular DNA repair functions, with implications for neuroblastoma treatment, risk stratification, and cancer predisposition.

## Introduction

High-risk neuroblastoma remains a significant clinical challenge, with mortality exceeding 50% despite intensive multimodal chemoradiotherapy and immune-based treatment regimens.^1^ In recent years, the genetic basis of neuroblastoma has come into focus. Germline variants in *ALK* are the predominant cause of familial neuroblastoma, accounting for 1-2% of overall cases,^2^ mutations in *PHOX2B* cause neuroblastoma in the context of a global neurocristopathy that can be familial,^3, 4^ and multiple genomic loci have been implicated in predisposition to the more common sporadic neuroblastoma through a large genome-wide association study (GWAS).^5, 6^ One of the most significantly neuroblastoma-associated genomic regions in this GWAS is a complex linkage disequilibrium (LD) block centered on the BRCA1-associated RING domain 1 (*BARD1*) gene.^7^ The BARD1 protein is a well-documented heterodimerization partner of BRCA1^8, 9^, a critical interaction required for BRCA1 stability, ubiquitin ligase activity, and other critical cellular functions including repair of DNA double-strand breaks (DSBs) by homologous recombination.^10–14^ Common single nucleotide polymorphisms (SNPs) at the *BARD1* locus are associated with neuroblastoma across multiple ethnicities and are enriched in high-risk patients.^5, 7, 15, 16^ Additionally, *BARD1* risk variants correlate with increased expression of an oncogenically activated BARD1 isoform and reduced expression of full-length *BARD1*.^7, 17–19^

More recently, rare coding variants with larger predicted effect sizes were also found to be significantly enriched in the germline of sporadic neuroblastoma patients, including multiple putative loss-of-function (LOF) variants in the *BARD1* gene.^20–22^ Notably, *BARD1* coding variants are also enriched in the germline of patients with several other malignancies^23–29^, suggesting a potential shared mechanism of tumor predisposition across multiple cancers. Large germline sequencing studies in pediatric and adult cancers have successfully described the landscape of cancer-associated germline variation across many malignancies^30–32^, but the precise functional implications of these germline variants on tumor development remain largely undefined. To address this need, here we evaluated the functional impact of both common and rare germline variation at the *BARD1* locus in neuroblastoma. Specifically, we aimed to determine whether *BARD1* variants perturb DNA repair efficiency in neuroblastoma cells.

## Brief Materials and Methods

Detailed methods are provided in the Supplemental Methods. *BARD1* isogenic cellular models were generated via CRISPR/Cas9 where Cas9 enzyme, guide RNAs and single-stranded mutated donor oligonucleotides were introduced into neuroblastoma cells via electroporation. Genotypes were confirmed by Sanger sequencing. Quantitative real-time (RT)-PCR results were derived via the 2ΔΔ^Ct^ method. Whole genome sequencing (WGS) was performed on isogenic and control cellular models. Quantification of DNA structural variants, copy number, indels, single-nucleotide variants (SNVs) and DSBs was performed as previously described.^33–37^

Immunofluorescence images were obtained using a Leica fluorescence microscope and standard staining protocols. RAD51 foci were quantified using Focinator v2.0 software and ImageJ.^38^ The mClover-LMNA assay was performed according to published methods.^39^ Cytotoxicity studies were performed via serial dilution of drugs and vehicle controls and cell viability was measured via CellTiter-Glo® assays*. In vivo* studies were performed as previously described^40^ using murine xenografts generated from *BARD1* isogenic cells.

## Results

### Neuroblastoma-associated *BARD1* common germline variation correlates with increased somatic DNA double strand breaks

We previously reported that common SNPs at the *BARD1* locus are significantly associated with neuroblastoma predisposition across multiple ethnicities^7, 15, 16, 18^ and that a subset of these SNPs are correlated with the expression of oncogenically activated BARD1 isoforms.^17^ However, we suspect that there are additional mechanisms involving *BARD1* that contribute to neuroblastoma predisposition. In view of BARD1’s critical role in stabilizing BRCA1 and facilitating accurate DNA repair^11, 41, 42^, we first tested the hypothesis that common *BARD1* variation is associated with reduced *BARD1* expression that results in a genome-wide DNA repair deficiency. We selected the most significant, directly genotyped common risk allele (T) at SNP rs17487792 from our GWAS^5^ for further analysis. We first queried the multi-tissue eQTL data from the Genotype-Tissue Expression portal (GTEx, https://gtexportal.org/) for the risk allele of this SNP and found that the T risk allele was associated with a significant reduction in *BARD1* expression across 34 normal tissues, notably including multiple nervous system tissues (**Figure 1A, B**; p = 1 x 10^-5^ - 3 x 10^-32^). Next, to determine if there was any association between the rs17487792 T risk allele and genomic instability in neuroblastoma tumors, we utilized two large sets of primary neuroblastoma tumors subjected to either paired tumor-normal WGS (n=134 tumors; cohort 1) or SNP genotyping (n=383 tumors; cohort 2). Using methods previously described,^35, 37^ we quantified the tumor burden of DNA double-strand breaks (DSBs) and correlated these data with rs17487792 genotype in these two tumor datasets. Tumors arising in children harboring a germline homozygous risk allele genotype (T/T) at SNP rs17487792 had a significantly increased burden of DSBs when compared to the homozygous non-risk allele genotype at this SNP (C/C; **Figure 1C-F**). This correlation between SNP rs17487792 genotype and quantity of DNA DSBs was more pronounced in patients harboring tumors without *MYCN* amplification, an association that was also found when limiting these analyses to only high-risk neuroblastomas (**Supplemental Figure 1A-F**). Taken together, these findings suggest that deficiencies in DNA repair associated with decreased *BARD1* expression are an additional mechanism by which common variants at the *BARD1* locus contribute to neuroblastoma predisposition.

**Figure 1.**
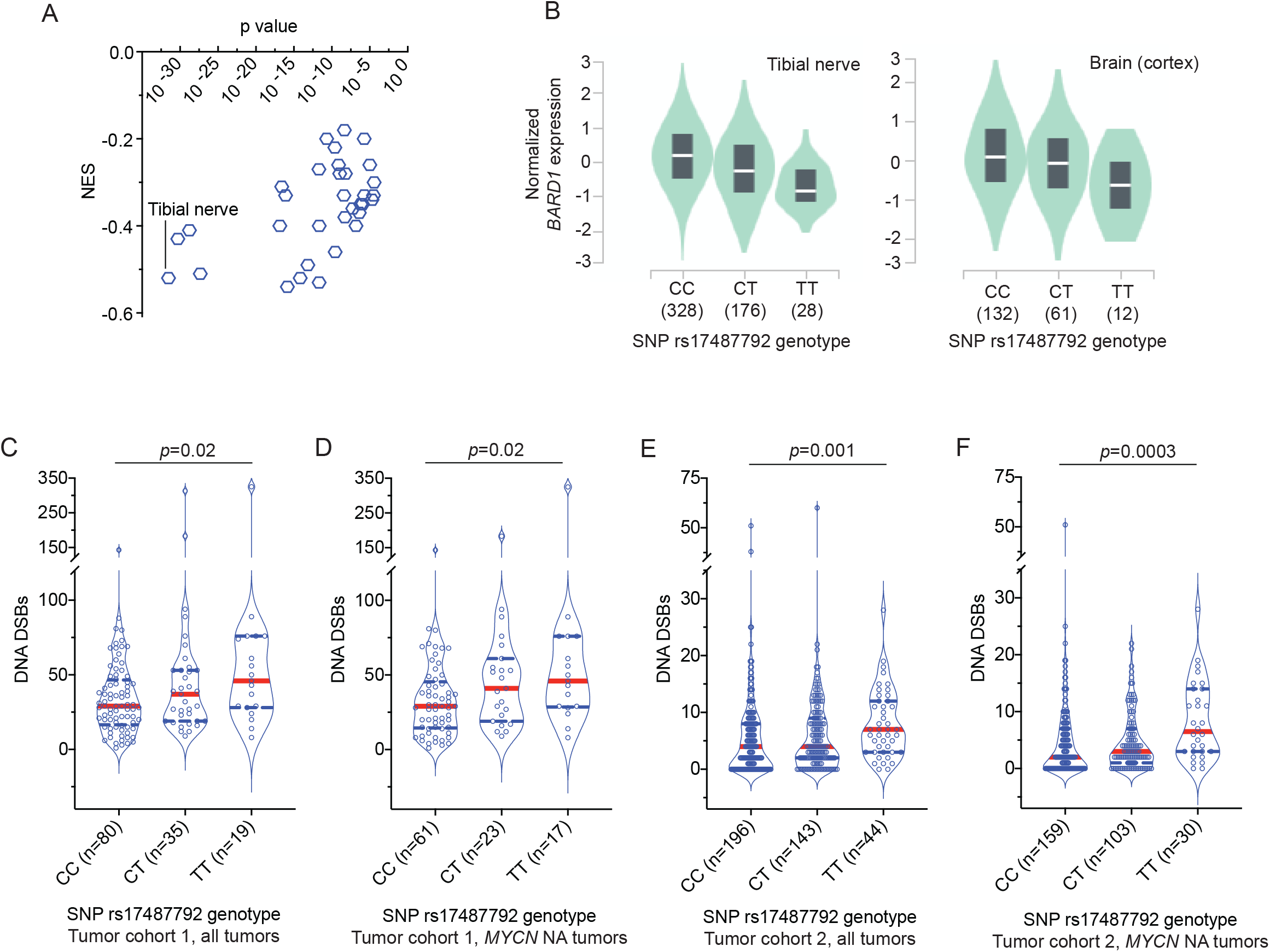
Common *BARD1* germline risk variants correlate with decreased *BARD1* expression and genome-wide deficiencies in DNA repair. (**A**) Plot of the normalized effect size (NES) of the SNP rs17487792 eQTL versus p value for 33 human tissues derived from the GTEx project. (**B**) Plots of *BARD1* expression versus SNP rs17487792 genotype in tibial nerve (left) and brain cortex (right) derived from the GTEx project. (**C-F**) Violin plots depicting the number of DNA DSBs in neuroblastoma tumors from patients with different germline SNP rs174877792 genotypes. Panels **C** and **D** depict quantity of DNA DSBs in cohort 1 among all tumors or among only tumors without *MYCN* amplification, respectively. Panels **E** and **F** depict quantity of DNA DSBs in cohort 2 among all tumors or among only tumors without *MYCN* amplification, respectively. Red dotted line denotes median and blue dotted lines denotes quartiles. *MYCN* NA, *MYCN* non-amplified.

### Generation of isogenic neuroblastoma cellular models harboring neuroblastoma-associated *BARD1* germline loss-of-function variants (*BARD1*^+/mut^)

In a parallel study of 786 patients with neuroblastoma, we observed that *BARD1* was the most significantly altered cancer predisposition gene in the germline of neuroblastoma patients.^22^ We identified rare pathogenic (P) or likely pathogenic (LP) nonsense, frameshift, or splice site variants in *BARD1* in the germline DNA in 8 of the 786 patients (1%).^22^ These variants were distributed throughout the *BARD1* coding sequence and displayed no evidence of somatic loss of heterozygosity (LOH) at the *BARD1* locus in available matched tumor DNA (**Supplemental Table 1**). Further, many of these variants are enriched in the germline DNA of adults with other malignancies (**Supplemental Table 1**).

We next aimed to study the functional implications of these germline P-LP loss of function (LOF) *BARD1* variants, focusing on DNA repair mechanisms of the BARD1-BRCA1 heterodimer.^11, 41, 42^ We introduced a subset of these *BARD1* variants as monoallelic knock-ins via CRISPR/Cas9 genome editing (**Supplemental Table 1** and **2**) utilizing two complementary cell lines to study their functional impact: IMR-5 [a *MYCN-* amplified, *TP53* wild-type (WT) neuroblastoma cell line; hereafter designated as IMR-5 *BARD1*^+/mut^] and hTERT RPE1 (an immortalized cell line of neural crest origin; hereafter designated as RPE1 *BARD1*^+/mut^).^43^ We chose to focus on the identified P-LP *BARD1* LOF nonsense and frameshift variants, rather than the *BARD1* splice site alterations or the common non-coding SNP variations identified via GWAS, hypothesizing that these LOF variants may result in the most significant and reproducible phenotypes. We successfully engineered four of these variants (*BARD1*^R112*, R150* E287fs and Q564*^) across these two cellular models (**Supplemental Table 1**). Heterozygosity for the appropriate *BARD1* variant was confirmed by Sanger sequencing (**Figure 2A**). The most likely exonic off-target CRISPR sites with cutting frequency determination (CFD) scores ≥ 0.04 (n = 2-3 loci per mutation) as determined by the CRISPOR tool^44^ were also sequenced to ensure no aberrant Cas9 editing had occurred (**Supplemental Table 3**). Clones with no evidence of editing at either *BARD1* allele, which had undergone similar single-cell selection pressure, were chosen at random to use as controls for subsequent sequencing and functional studies.

**Figure 2.**
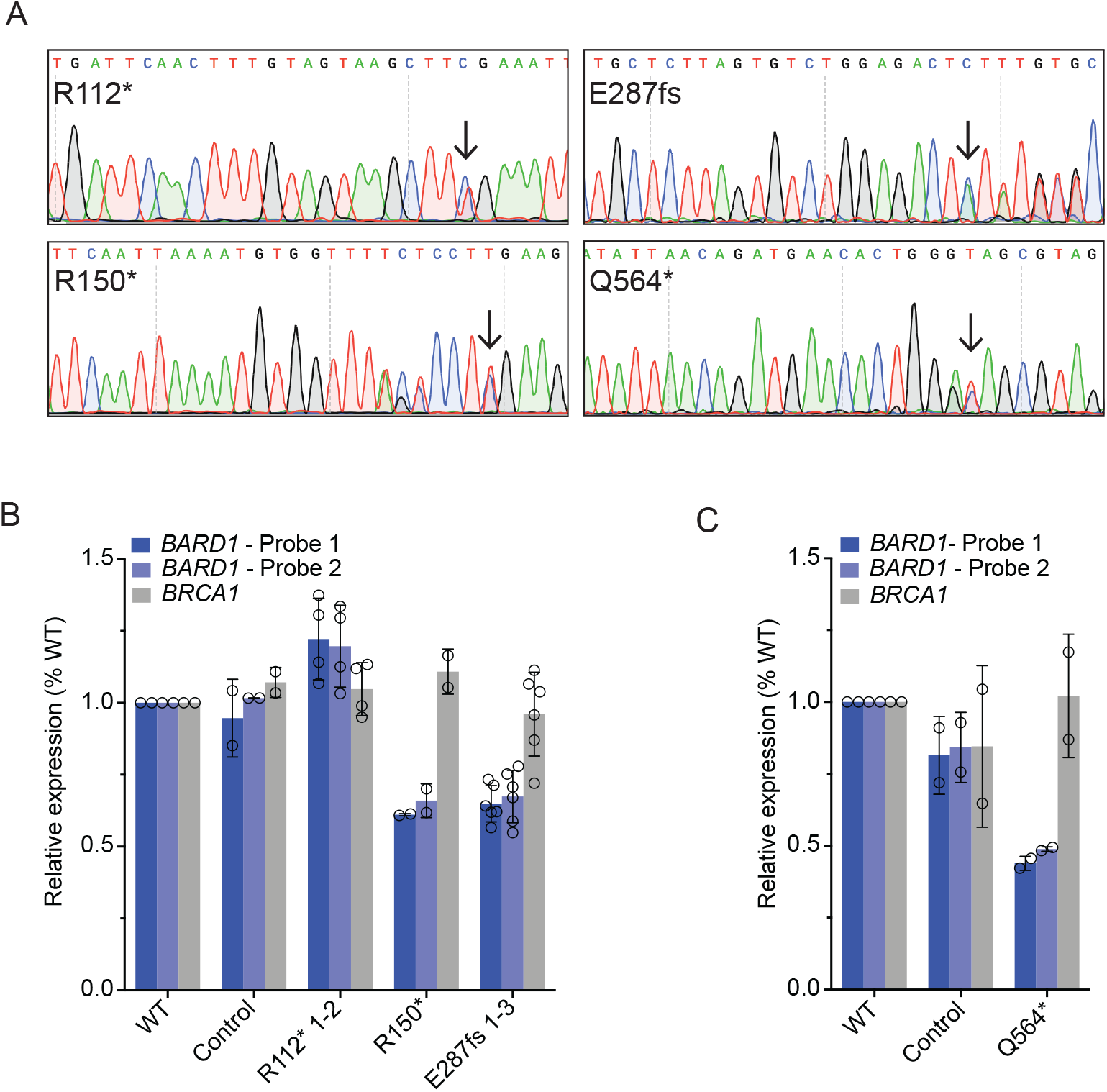
Neuroblastoma cells heterozygous for disease-associated *BARD1* loss-of-function variants (*BARD1*^+/mut^) have reduced *BARD1* expression. (**A**) Representative chromatograms from IMR-5 and RPE1 *BARD1*^+/mut^ isogenic cell lines. Black arrows indicate CRISPR-introduced *BARD1* heterozygous variants. Other variants reflect synonymous PAM changes (R150*, Q564*) or frameshift-induced nucleotide alterations (E287fs). (**B, C**) *BARD1* and *BRCA1* expression in IMR-5 *BARD1*^+/mut^ (**B**) and RPE1 *BARD1*^+/mut^ (**C**) cells and nontargeted clonal control cells. *BARD1* expression measured via two unique TaqMan® probes. **B** and **C** are represented as means ± SD of 2 biological replicates of each unique cell line, including multiple cell lines with identical *BARD1* variants (n = 2 IMR-5 *BARD1*^+/R112*^, n = 1 for IMR-5 *BARD1*^+/R150*^ and n = 3 for IMR-5 *BARD1*^+/E287fs^ cell lines).

### Cells heterozygous for *BARD1* loss-of-function variants have reduced *BARD1* expression

We first sought to characterize *BARD1* expression in these IMR-5 and RPE1 *BARD1*^+/mut^ cellular models, given that similar *BRCA1* monoallelic variants induce functionally relevant *BRCA1* haploinsufficiency^45, 46^ and considering our finding that neuroblastoma-associated common *BARD1* variation is significantly associated with decreased *BARD1* expression. Via RT-PCR using two unique *BARD1* TaqMan® probes, we found that three of four *BARD1*^+/mut^ isogenic cellular models exhibited a substantial reduction in *BARD1* expression compared to WT parental cells or non-targeted clonal control cells (33-56% reduction in *BARD1* mRNA in IMR-5/RPE1 *BARD1*^+/mut^ cells; **Figure 2B, C**). However, IMR-5 *BARD1*^+/R112*^ cells had *BARD1* mRNA expression comparable to WT cells and non-targeted clonal control cells (**Figure 2B**). The *BARD1*^R112*^ variant is unique among the prioritized *BARD1* nonsense and frameshift variants studied in that it has a nearby downstream putative start codon (M145) distal to the aberrant stop codon, potentially allowing for resumption of translation and avoidance of nonsense-mediated decay, a phenomenon that has been previously described.^47^ We also quantified *BRCA1* mRNA as a control and found no difference in *BRCA1* mRNA expression between WT, clonal control cells, and *BARD1*^+/mut^ isogenic cells (**Figure 2B, C**). Thus, most of the prioritized neuroblastoma associated *BARD1* LOF germline variants result in reduced *BARD1* expression.

### *BARD1*^+/mut^ cells exhibit widespread genomic instability

The BRCA1-BARD1 heterodimer is essential for maintaining genomic integrity, in part via the homology-directed repair (HDR) of DNA DSBs, and significant HDR defects have been observed in cellular models with heterozygous LOF *BRCA1* mutants.^10, 45, 48^ Considering these data, along with the correlation of neuroblastoma-associated common variation with deficient DNA repair, we first quantified the genome-wide impact of this *BARD1* haploinsufficiency on DNA repair in isogenic IMR-5 *BARD1*^+/mut^ cells. After twenty passages in cell culture, genomic DNA from three representative IMR-5 *BARD1*^+/mut^ clones, a non-targeted control clone, and WT parental cells were examined via whole-genome sequencing. We identified genomic aberrations that each clonal cell line acquired relative to the WT parental cell line by applying Control-FREEC^33^ and Delly^34^ as complementary algorithms for copy number and structural variant analysis, respectively. Striking genomic instability was observed uniquely in the IMR-5 *BARD1*^+/mut^ isogenic cell lines, including large-scale copy number alterations (**Figure 3A, Supplemental Figure 2A**) and structural variants (**Figure 3B-C, Supplemental Figure 2B-C**). We also quantified DNA DSBs^35, 37^ based on both Control-FREEC and Delly calls, and all three IMR-5 *BARD1*^+/mut^ isogenic cell lines harbored substantially increased DSBs compared to the non-targeted control clone (**Figure 3D**). These results were consistent using both stringent (**Figure 3B-D, Supplemental Figure 2B-C**) and more relaxed structural variant filtering parameters (**Supplemental Figure 3A-E**). Parallel analyses with MuTect^36^ also revealed an increase in SNVs and indels in IMR-5 *BARD1*^+/mut^ clones versus the control clone, with variants in *BARD1*^+/mut^ clones exhibiting higher allele frequencies (**Supplemental Figure 4A-C**). Finally, analysis of mutational signatures in the non-targeted control and isogenic IMR-5 *BARD1*^+/mut^ cell lines revealed enhanced exposure of SBS3 (Defective HR DNA repair; BRCA1/2 mutation) in the isogenic cell lines, along with other DNA repair deficiency signatures (SBS6, Defective DNA mismatch repair; SBS10, POLE mutation) uniquely found in a subset of the isogenic IMR-5 *BARD1*^+/mut^ cell lines but not the non-targeted control cells (**Figure 3E, Supplemental Figure 4D**).

**Figure 3.**
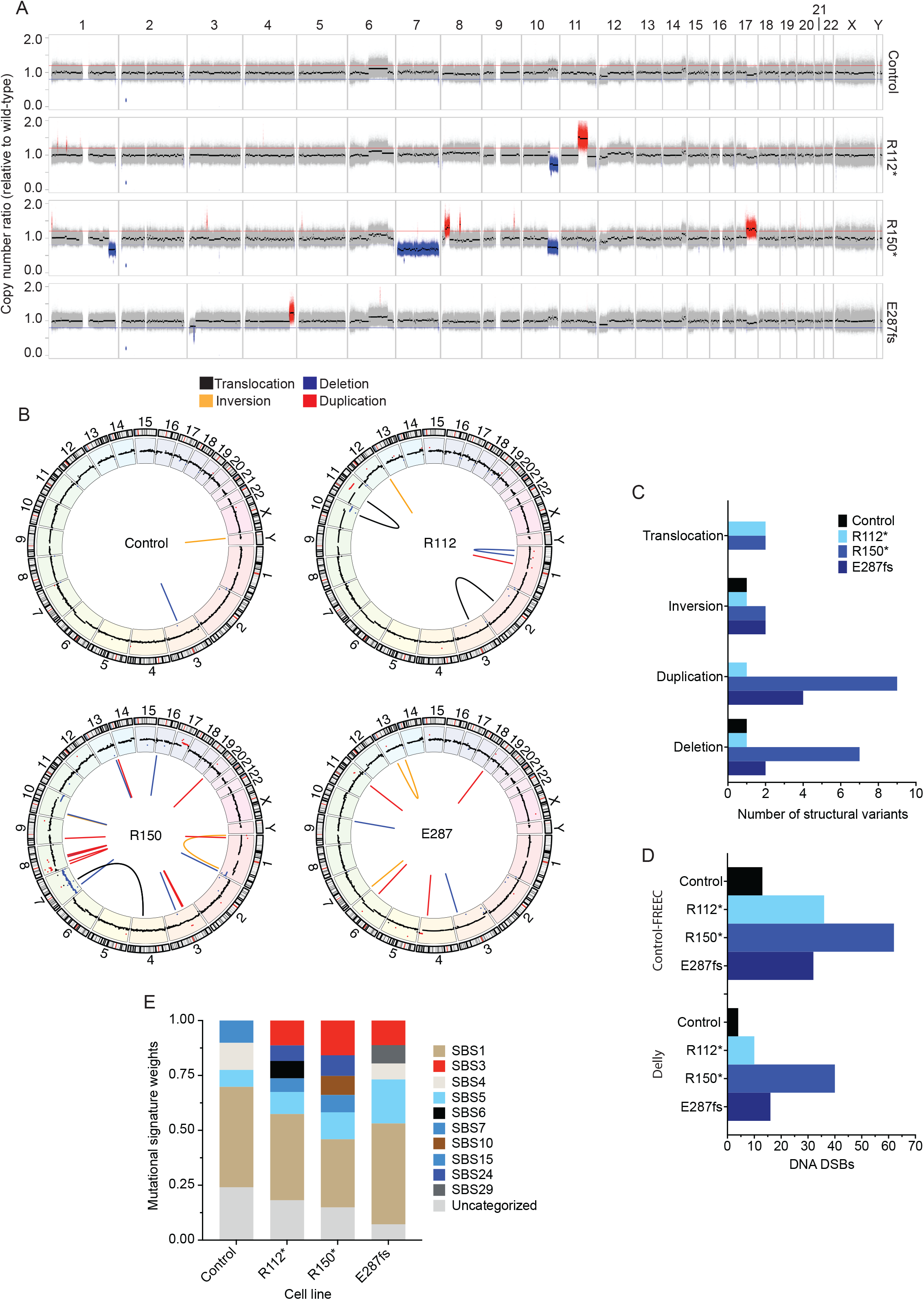
Neuroblastoma IMR-5 *BARD1*^+/mut^ cell lines exhibit widespread genomic instability. (**A**) Copy number ratio across the genome for the IMR-5 non-targeted control clone and *BARD1*^+/mut^ isogenic clones relative to WT parental IMR-5 cells. Large acquired genomic losses are indicated in blue (fold change < 0.8) and gains in red (fold change > 1.2). (**B**) Circos plots depicting structural variants identified in non-targeted control and *BARD1*^+/mut^ IMR-5 cells. Copy number segments from (**A**) are shown in the outer circle for reference. (**C**) Counts of structural variants in non-targeted control and *BARD1*^+/mut^ IMR-5 cells. (**D**) Counts of DNA DSBs in non-targeted control and *BARD1*^+/mut^ IMR-5 cells, quantified from the Control-FREEC copy number data (**top**) and Delly structural variant data (**bottom**). (**E**) Plot of mutational signature weights in non-targeted control and *BARD1*^+/mut^ IMR-5 cells using COSMIC mutational signatures (v2).

### *BARD1*^+/mut^ cells are deficient in DNA repair and are more sensitive to cisplatin and PARP inhibition

We next used several complementary functional studies to evaluate the ability of *BARD1*^+/mut^ isogenic cells to perform efficient DNA repair. First, we utilized a CRISPR-based *in vitro* assay incorporating a fluorescent endogenous mClover tag (**Figure 4A**) to directly quantify HDR efficiency in the *BARD1*^+/mut^ isogenic models via flow cytometry.^39^ We found that after Cas9-induced DNA cutting, IMR-5 *BARD1*^+/mut^ isogenic cellular models consistently integrated the DNA repair template with the mClover tag at approximately 50% the efficiency of WT IMR-5 cells (relative mean mClover-positive *BARD1*^+/mut^ isogenic cells 46-53% of WT IMR-5 cells; **Figure 4B-C**). Notably, the relative reduction in HDR capacity was consistent among the different isogenic cell lines, except one of the three IMR-5^+/E287fs^ isogenic cell clones which had similar levels of mClover tag integration as WT IMR-5 cells (blue triangles; **Figure 4C**).

**Figure 4.**
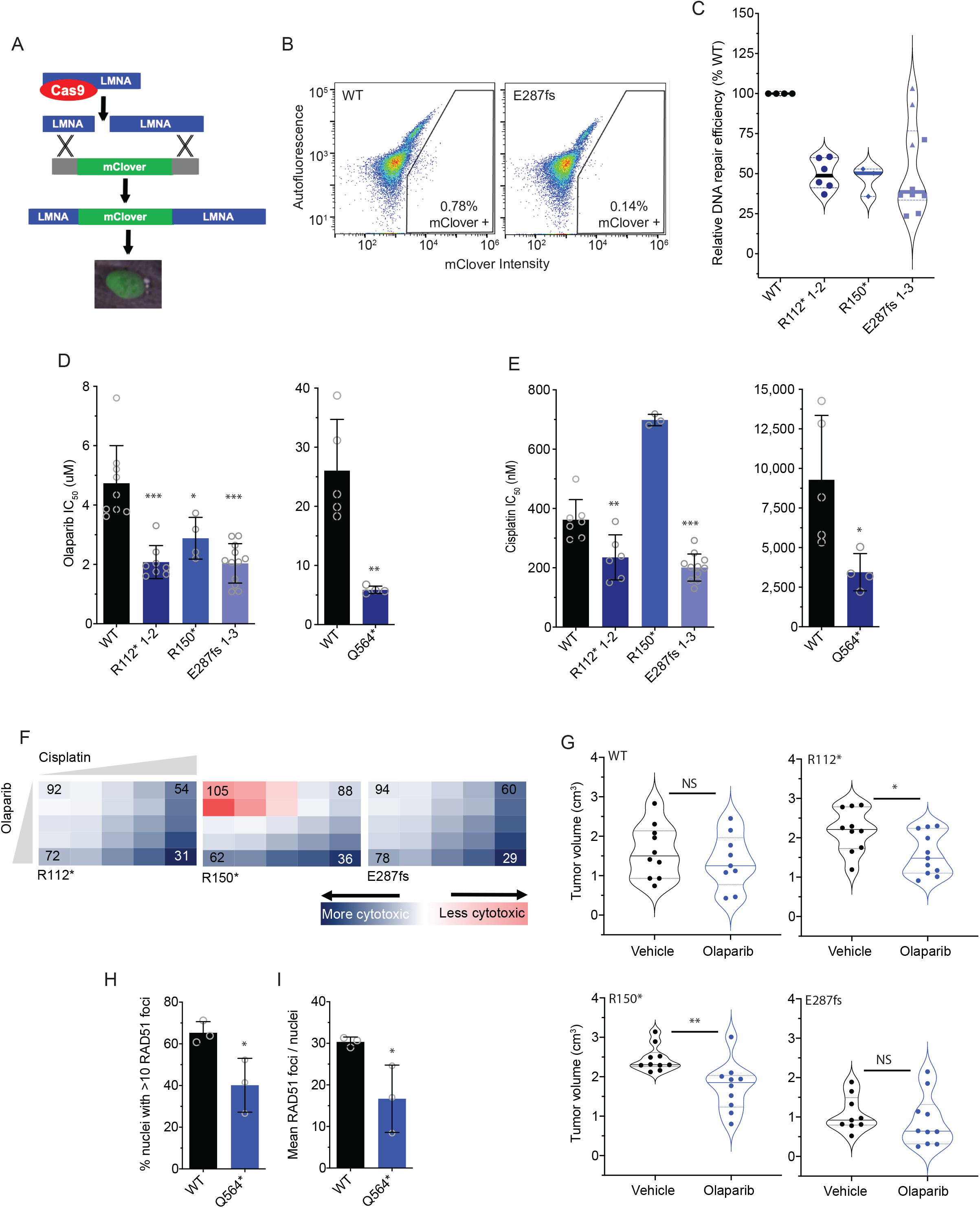
*BARD1*^+/mut^ cells are deficient in DNA repair and sensitive to PARP inhibition. (**A**) Schematic of the mClover-LMNA HDR assay.^39^ (**B**) Representative flow cytometry plots of IMR-5 WT and *BARD1*^+/E287fs^ cells co-transfected with pX330-LMNA1 gRNA and pCR2.1-CloverLamin repair template plasmids with gating strategy for clover-positive cells indicated. (**C**) Violin plots showing relative DNA damage repair efficiency across IMR-5 *BARD1*^+/mut^ cells as quantified with the mClover-LMNA HDR assay shown in **A** and **B**. (**D**) Olaparib IC50 values in IMR-5 WT and *BARD1*^+/mut^ cell lines (**left**) and in RPE1 WT and *BARD1*^+/mut^ cell lines (**right**). (**E**) Cisplatin IC50 values in IMR-5 WT and *BARD1*^+/mut^ cell lines (**left**) and in RPE1 WT and *BARD1*^+/mut^ cell lines (**right**). (**F**) Relative cytotoxicity of combined olaparib and cisplatin in IMR-5 *BARD1*^+/mut^ cell lines compared to WT. Each square represents a single dose combination. Blue squares represent drug combinations at which greater cytotoxicity is observed in *BARD1*^+/mut^ cells; red squares represent drug combinations at which greater cytotoxicity is observed in WT cells. Numbers in corner cells represent the percent of isogenic cells alive compared to WT cells alive at equivalent doses. (**G**) Violin plots of tumor volumes after 2 weeks of olaparib treatment in *BARD1*^+/mut^ versus WT IMR-5 xenografts. (n=9-11 mice/cohort). Tumor volumes for olaparib-treated BARD1 E287fs xenografts measured on day 13. Solid lines denote medians and dotted lines denote quartiles. (**H**) Proportion of RPE1 WT and RPE1^+/Q564*^ nuclei with >10 RAD51 foci after treatment with cisplatin. (**I**) Mean RAD51 foci per RPE1 WT and RPE1^+/Q564*^ nucleus after treatment with cisplatin. Data in **C-E** are means ± SD of 3-12 biological replicates of each isogenic cell line, including multiple cell lines with identical *BARD1* variants (n = 2 IMR-5 *BARD1*^+/R112*^, n = 1 for IMR-5 *BARD1*^+/R150*^ and n = 3 for IMR-5 *BARD1*^+/E287fs^ cell lines). LMNA, lamin A/C; *p < 0.05, **p < 0.01, ***p < 0.0001; NS, not significant as measured by T test.

We next investigated whether *BARD1*^+/mut^ cells displayed increased vulnerability to PARP inhibition, another well-validated marker of DNA damage repair deficiency.^49^ We treated both WT and isogenic IMR-5 *BARD1*^+/mut^ cells with the PARP inhibitor olaparib (0.1 - 100 μM) and assessed cytotoxicity after four days (**Figure 4D**). IMR-5 *BARD1*^+/mut^ cells exhibited significantly enhanced sensitivity to olaparib compared to WT cells (mean IMR-5^+/mut^ IC50 2.1 - 2.9 μM versus 4.7 μM for WT IMR-5 cells; *P* < 0.05, **Figure 4D, left**), as did RPE1 *BARD1*^+/Q564*^ cells (mean IC50 5.9 μM versus 26.0 μM for WT RPE1 cells; *P* < 0.01, **Figure 4D, right**). Similarly, except for IMR-5 *BARD1*^/R150+*^, *BARD1*^+/mut^ cells also displayed increased sensitivity to the DNA intercalating agent cisplatin [mean IC_50_ 201-235 nM for IMR-5 *BARD1*^+/mut^ cells versus 362 nM for WT IMR-5 cells (*P* < 0.01; **Figure 4E, left**) and 3,450 nM for RPE1 *BARD1*^+/mut^ cells versus 9,282 nM for WT RPE1 cells; *P* < 0.05; **Figure 4E, right**]. To further validate these data, we designed a cytotoxicity assay incorporating both olaparib and cisplatin at escalating doses to evaluate whether the combination was also more potent in IMR-5 *BARD1*^+/mut^ cells than WT IMR-5 cells. Compared with WT IMR-5 cells, *BARD1*^+/R112*^ and *BARD1*^+/E287fs^ IMR-5 cells were more sensitive to these drugs at every dose combination tested, and *BARD1*^+/R150*^ cells were more sensitive at most doses (**Figure 4F**). Next, to confirm the sensitivity of *BARD1*^+/mut^ cells to olaparib, we treated cohorts of mice xenografted with each isogenic line with 20 mg/kg of olaparib daily for 28 days.^50^ Olaparib-treated *BARD1*^+/R112*^ and *BARD1*^+/R150*^ isogenic xenografts had a significant reduction in tumor growth versus paired vehicle-treated animals, while WT IMR-5 and the much slower-growing IMR-5 *BARD1*^+/E287fs^ xenografts did not show any olaparib-induced tumor growth delay (**Figure 4G, Supplemental Figure 5A-D**). Mice harboring the *BARD1*^/R150*^ xenografts also had significantly longer event-free survival compared to paired-vehicle treated mice (*P* < 0.01; **Supplemental Figure 5E-H**).

Finally, we utilized the RPE1 isogenic model to evaluate the ability of *BARD1*^+/mut^ cells to form RAD51 foci after treatment with cisplatin.^51^ The larger nucleus of RPE1 cells facilitated increased resolution of distinct RAD51 foci by immunofluorescence. Twenty-four hours after treatment with cisplatin, significantly fewer RPE1 *BARD1*^+/Q564*^ cells exhibited >10 RAD51 foci per nucleus (mean of 40.1% versus 65.3% for WT RPE1 cells, *P* < 0.05; **Figure 4H**), and nuclei from RPE1 *BARD1*^+/Q564*^ cells had significantly fewer average RAD51 foci per nucleus overall (mean of 17 versus 30 for RPE1 WT cells; *P* < 0.05, **Figure 4I**). Taken together, these findings suggest that neuroblastoma-associated *BARD1* LOF variants impair DNA repair functions.

## Discussion

Neuroblastoma, like all human cancers, is a genetic disease. Common germline alleles at several genomic loci (*e.g., BARD1, LMO1, CASC15*) contribute to sporadic neuroblastoma predisposition, and rare heterozygous variants in neurodevelopmental genes (*PHOX2B* and *ALK*) underlie familial neuroblastoma.^2–4, 20^ More recently, next-generation sequencing efforts focused on germline DNA from neuroblastoma patients have also identified enrichment of potentially pathogenic variants in cancer predisposition genes; among these, the most frequently altered gene is *BARD1.^22^* While significant progress has been made in defining the landscape of rare germline variation in cancer predisposition genes across pediatric and adult cancers^30–32^, less effort has concentrated on elucidating how these disease-associated genetic variants influence cancer development at the molecular level, especially in pediatric malignancies. Thus, the functional validation of neuroblastoma-associated *BARD1* germline variants described here not only further enhances our understanding of the contribution of the *BARD1* locus to neuroblastoma predisposition, but also represents a critical attempt to define the functional and potential clinical relevance of cancer-associated germline variation.

Given that *BARD1* is one of the most significant and replicated neuroblastoma-associated GWAS loci, we first looked for genome-wide evidence of a DNA damage repair defect in tumors from patients harboring a common *BARD1* risk haplotype. We quantified DNA DSBs in two large sets of neuroblastoma tumors and observed a correlation between germline SNP genotype and somatic DNA damage, which was most robust in tumors without *MYCN* amplification. Given MYCN’s central role in response to DNA damage^52, 53^, this suggests a potential compensatory mechanism for DNA repair in neuroblastoma cells with high levels of MYCN. However, given the low effect sizes of these neuroblastoma-associated common variants, we chose to primarily focus on functional validation of a subset of the recently identified LOF *BARD1* germline variants with much larger predicted effect sizes. Our findings support a model in which *BARD1* LOF variants induce *BARD1* haploinsufficiency leading to genomic instability from deficient DNA damage repair. This model may be immediately relevant to other predicted LOF variants in DNA repair-related genes that are enriched in the germline of children (and adults) with multiple tumor histotypes.^30, 32, 54^

In considering this haploinsufficiency model for heterozygous *BARD1* LOF variants in neuroblastoma tumorigenesis, it is important to note that the traditional model for *BRCA1-* and *BRCA2*-associated familial malignancies involves early LOH of the WT allele.^55^ Further, while emerging evidence suggests that a subset of breast and ovarian tumors from individuals with *BRCA1* or *BRCA2* germline variants lack *BRCA1/2* locus-specific LOH, these heterozygous tumors exhibit homologous recombination (HR) deficiency scores similar to non-*BRCA1/2*-mutated tumors.^56^ In contrast, a single hit to an HR pathway gene may be sufficient to induce HR deficiency in neuroblastoma and other pediatric cancers as observed in a recent pan-pediatric cancer study^57^ that identified eight patients with monoallelic germline variants in HR pathway genes such as *BRCA2.* Although none of the matched tumors in this study carried a second hit to induce locus-specific LOH, they exhibited mutational signatures consistent with HR deficiency. Additionally, considering that homozygous loss of either *BARD1* or *BRCA1* results in embryonic lethality in murine models,^58^ LOH may not be well-tolerated as an early event in pediatric cancers that commonly initiate during embryogenesis.

As germline sequencing of pediatric cancer patients becomes more widely adopted into clinical practice, the role of germline variants in treatment stratification, normal tissue susceptibility to cancer therapies, and implications for genetic counseling must all be rigorously examined. For example, here we suggest tumors harboring *BARD1* LOF variants may be more sensitive to DNA damaging agents, findings that may have important implications for selection of treatment regimens especially in the relapse setting where several different chemotherapeutic agents may be available to patients. Additionally, the presence of cancer-associated gene variants within normal tissues may result in significant morbidities from cytotoxic therapies. This phenomenon has been noted in cancer predisposition syndromes such as Li-Fraumeni, in which genotoxic treatments can precipitate severe acute toxicity and contribute to an elevated risk of second malignancy.^59, 60^ Additionally, of note during the preparation of this manuscript, a patient presented to our oncology clinic with a rare composite neuroblastoma/pheochromocytoma tumor that was ultimately found to have a *BARD1*^R150*^ germline variant. This patient was treated with our standard high-risk neuroblastoma regimen, including conditioning with carboplatin/etoposide/melphalan (CEM) followed by autologous hematopoietic stem cell transplantation,^61^ which precipitated severe acute kidney injury progressing rapidly to renal failure and death. While the contribution of this patient’s germline *BARD1* variant to this fatal complication cannot be known, this case raises the possibility that the *BARD1*^R150*^ variant caused increased renal sensitivity to the carboplatin-containing transplant conditioning regimen. A prior report from our institution focused on the toxicities in 44 patients who received the same CEM conditioning places this patient’s complication in context.^62^ Approximately one-third of these patients developed modest increases in serum creatinine, with two requiring brief courses of dialysis, but none experienced irreversible or fatal kidney injury. Clearly, the influence of germline variation in DNA repair-related genes, or other genes critical in normal tissue homeostasis, on the development of normal tissue toxicity during multimodal cancer therapy merits further investigation. Finally, the identification of pathogenic variants in the germline of children with cancer has important genetic counseling implications as a majority of these are likely inherited in an autosomal manner similar to recent findings in pediatric sarcomas.^63^ While these moderate penetrance pathogenic variants are unlikely to be sufficient to cause malignancy alone, they do substantially increase the risk of developing cancer, which has important repercussions for family members who may also harbor an identical germline genotype. Furthermore, it is now clear that pathogenic germline variants classically associated with adult cancer predisposition syndromes (*e.g., BRCA1/2*) also contribute to cancer risk in children and adolescents,^64^ further emphasizing the importance for cascade testing.

Lastly, while this study focused specifically on germline loss-of-function variants, other heterozygous germline and somatic variations in *BARD1* have also been identified.^20, 22^ These variants may induce similar defects in DNA repair efficiency and genomic stability. The functional validation approach taken here can be extended to these and other potentially pathogenic germline variants recently identified across several childhood and adult malignancies, especially those variants that are predicted to disrupt DNA repair pathways. Finally, while at some larger academic medical centers it is common practice to sequence both tumor and germline tissues at diagnosis, these data support adopting this parallel sequencing practice for all pediatric cancer patients. Future studies will be essential to both illuminate additional functional implications of cancer-predisposing germline genetic variants in tumorigenesis and to expand their utility in oncology clinical practice.

## Acknowledgements

This research was supported by a Howard Hughes Medical Institute Medical Fellows grant (M.P.R.). K.R.B. is a Damon Runyon Physician-Scientist supported (in part) by the Damon Runyon Cancer Research Foundation (PST-07-16). L.E.E. was supported in part by NIH T32 GM008216. This work was also supported by an Alex’s Lemonade Stand Young Investigator Award (K.R.B.), the EVAN Foundation (K.R.B.), the Giulio D’Angio Endowed Chair (J.M.M.), NCI K08 CA230223 (K.R.B.), NCI R35 CA220500 (J.M.M.), NCI R01 CA204974 (S.J.D.), NCI R01 CA237562 (S.J.D.), an Early Researcher Award from the Ontario Ministry of Research and Innovation (A.S.), the Canada Research Chair in Childhood Cancer Genomics (A.S.), the V Foundation (A.S.), and the Robert J. Arceci Innovation Award from the St. Baldrick’s Foundation (A.S.).

## Author Contributions

Conceptualization, M.P.R., L.E.E., J.M.M., S.J.D. and K.R.B. Methodology, M.P.R., L.E.E., Z.V., S.J.D. and K.R.B, Experimental Investigation, M.P.R., M.S., M.T., D.G. and K.R.B. Computational Investigation, L.E.E., Z.V., J.P.E., J.L.R., J.W., M.L., A.S., and S.J.D. Funding Acquisition, M.P.R., J.M.M., S.J.D. and K.R.B. Writing, M.P.R., L.E.E. and K.R.B.

## References

1. Maris JM. Recent advances in neuroblastoma. N Engl J Med. Jun 10 2010;362(23):2202–11. doi:10.1056/NEJMra0804577

2. Mosse YP, Laudenslager M, Longo L, et al. Identification of ALK as a major familial neuroblastoma predisposition gene. Nature. Oct 16 2008;455(7215):930–5. doi:10.1038/nature07261

3. Trochet D, Bourdeaut F, Janoueix-Lerosey I, et al. Germline Mutations of the Paired-Like Homeobox 2B (PHOX2B) Gene in Neuroblastoma. Am J Hum Genet. Apr 2004;74(4):761–4.

4. Mosse YP, Laudenslager M, Khazi D, et al. Germline PHOX2B Mutation in Hereditary Neuroblastoma. Am J Hum Genet. Oct 2004;75(4):727–30.

5. McDaniel LD, Conkrite KL, Chang X, et al. Common variants upstream of MLF1 at 3q25 and within CPZ at 4p16 associated with neuroblastoma. PLoS Genet. May 2017;13(5):e1006787. doi:10.1371/journal.pgen.1006787

6. Maris JM, Mosse YP, Bradfield JP, et al. Chromosome 6p22 locus associated with clinically aggressive neuroblastoma. N Engl J Med. Jun 12 2008;358(24):2585–93. doi:10.1056/NEJMoa0708698

7. Capasso M, Devoto M, Hou C, et al. Common variations in BARD1 influence susceptibility to high-risk neuroblastoma. Nat Genet. Jun 2009;41(6):718–23. doi:10.1038/ng.374

8. Baer R, Ludwig T. The BRCA1/BARD1 heterodimer, a tumor suppressor complex with ubiquitin E3 ligase activity. Curr Opin Genet Dev. Feb 2002;12(1):86–91. doi:10.1016/s0959-437x(01)00269-6

9. Brzovic PS, Rajagopal P, Hoyt DW, King MC, Klevit RE. Structure of a BRCA1-BARD1 heterodimeric RING-RING complex. Nat Struct Biol. Oct 2001;8(10):833–7. doi:10.1038/nsb1001-833

10. Zhao W, Steinfeld JB, Liang F, et al. BRCA1-BARD1 promotes RAD51-mediated homologous DNA pairing. Nature. Oct 19 2017;550(7676):360–365. doi:10.1038/nature24060

11. Simons AM, Horwitz AA, Starita LM, et al. BRCA1 DNA-binding activity is stimulated by BARD1. Cancer Res. Feb 15 2006;66(4):2012–8. doi:10.1158/0008-5472.CAN-05-3296

12. Fabbro M, Savage K, Hobson K, et al. BRCA1-BARD1 complexes are required for p53Ser-15 phosphorylation and a G1/S arrest following ionizing radiation-induced DNA damage. J Biol Chem. Jul 23 2004;279(30):31251–8. doi:10.1074/jbc.M405372200

13. Joukov V, Groen AC, Prokhorova T, et al. The BRCA1/BARD1 heterodimer modulates ran-dependent mitotic spindle assembly. Cell. Nov 3 2006;127(3):539–52. doi:10.1016/j.cell.2006.08.053

14. Starita LM, Horwitz AA, Keogh MC, Ishioka C, Parvin JD, Chiba N. BRCA1/BARD1 ubiquitinate phosphorylated RNA polymerase II. J Biol Chem. Jul 1 2005;280(26):24498–505. doi:10.1074/jbc.M414020200

15. Capasso M, Diskin SJ, Totaro F, et al. Replication of GWAS-identified neuroblastoma risk loci strengthens the role of BARD1 and affirms the cumulative effect of genetic variations on disease susceptibility. Research Support, Non-U.S. Gov’t. Carcinogenesis. Mar 2013;34(3):605–11. doi:10.1093/carcin/bgs380

16. Latorre V, Diskin SJ, Diamond MA, et al. Replication of neuroblastoma SNP association at the BARD1 locus in African-Americans. Cancer Epidemiol Biomarkers Prev. Apr 2012;21(4):658–63. doi:10.1158/1055-9965.EPI-11-0830

17. Bosse KR, Diskin SJ, Cole KA, et al. Common variation at BARD1 results in the expression of an oncogenic isoform that influences neuroblastoma susceptibility and oncogenicity. Cancer Res. Apr 15 2012;72(8):2068–78. doi:10.1158/0008-5472.CAN-11-3703

18. Cimmino F, Avitabile M, Diskin SJ, et al. Fine mapping of 2q35 high-risk neuroblastoma locus reveals independent functional risk variants and suggests full-length BARD1 as tumor-suppressor. Int J Cancer. Dec 1 2018;143(11):2828–2837. doi:10.1002/ijc.31822

19. Cimmino F, Avitabile M, Lasorsa VA, et al. Functional characterization of full-length BARD1 strengthens its role as a tumor suppressor in neuroblastoma. J Cancer. 2020;11(6):1495–1504. doi:10.7150/jca.36164

20. Pugh TJ, Morozova O, Attiyeh EF, et al. The genetic landscape of high-risk neuroblastoma. Nat Genet. Mar 2013;45(3):279–84. doi:10.1038/ng.2529

21. Lasorsa VA, Formicola D, Pignataro P, et al. Exome and deep sequencing of clinically aggressive neuroblastoma reveal somatic mutations that affect key pathways involved in cancer progression. Oncotarget. Apr 19 2016;7(16):21840–52. doi:10.18632/oncotarget.8187

22. Kim J, Vaksman V, Egolf LE, et al. Germline pathogenic variants in cancer predisposition genes associate with worse survival and implicate BARD1 and DNA repair defects in neuroblastoma. JAMA Oncology, submitted, 2022.

23. Weber-Lassalle N, Borde J, Weber-Lassalle K, et al. Germline loss-of-function variants in the BARD1 gene are associated with early-onset familial breast cancer but not ovarian cancer. Breast Cancer Res. Apr 29 2019;21(1):55. doi:10.1186/s13058-019-1137-9

24. Venier RE, Maurer LM, Kessler EM, et al. A germline BARD1 mutation in a patient with Ewing Sarcoma: Implications for familial testing and counseling. Pediatr Blood Cancer. Sep 2019;66(9):e27824. doi:10.1002/pbc.27824

25. DeLeonardis K, Sedgwick K, Voznesensky O, et al. Challenges in Interpreting Germline Mutations in BARD1 and ATM in Breast and Ovarian Cancer Patients. Breast J. Jul 2017;23(4):461–464. doi:10.1111/tbj.12764

26. De Brakeleer S, De Greve J, Desmedt C, et al. Frequent incidence of BARD1-truncating mutations in germline DNA from triple-negative breast cancer patients. Clin Genet. Mar 2016;89(3):336–40. doi:10.1111/cge.12620

27. Ramus SJ, Song H, Dicks E, et al. Germline Mutations in the BRIP1, BARD1, PALB2, and NBN Genes in Women With Ovarian Cancer. J Natl Cancer Inst. Nov 2015;107(11) doi:10.1093/jnci/djv214

28. Schulz E, Valentin A, Ulz P, et al. Germline mutations in the DNA damage response genes BRCA1, BRCA2, BARD1 and TP53 in patients with therapy related myeloid neoplasms. J Med Genet. Jul 2012;49(7):422–8. doi:10.1136/jmedgenet-2011-100674

29. Ghimenti C, Sensi E, Presciuttini S, et al. Germline mutations of the BRCA1-associated ring domain (BARD1) gene in breast and breast/ovarian families negative for BRCA1 and BRCA2 alterations. Genes Chromosomes Cancer. Mar 2002;33(3):235–42. doi:10.1002/gcc.1223

30. Grobner SN, Worst BC, Weischenfeldt J, et al. The landscape of genomic alterations across childhood cancers. Nature. Mar 15 2018;555(7696):321–327. doi:10.1038/nature25480

31. Huang KL, Mashl RJ, Wu Y, et al. Pathogenic Germline Variants in 10,389 Adult Cancers. Cell. Apr 5 2018;173(2):355–370 e14. doi:10.1016/j.cell.2018.03.039

32. Zhang J, Walsh MF, Wu G, et al. Germline Mutations in Predisposition Genes in Pediatric Cancer. N Engl J Med. Dec 10 2015;373(24):2336–2346. doi:10.1056/NEJMoa1508054

33. Boeva V, Popova T, Bleakley K, et al. Control-FREEC: a tool for assessing copy number and allelic content using next-generation sequencing data. Bioinformatics. Feb 1 2012;28(3):423–5. doi:10.1093/bioinformatics/btr670

34. Rausch T, Zichner T, Schlattl A, Stutz AM, Benes V, Korbel JO. DELLY: structural variant discovery by integrated paired-end and split-read analysis. Bioinformatics. Sep 15 2012;28(18):i333–i339. doi:10.1093/bioinformatics/bts378

35. Lopez G, Conkrite KL, Doepner M, et al. Somatic structural variation targets neurodevelopmental genes and identifies SHANK2 as a tumor suppressor in neuroblastoma. Genome Res. Sep 2020;30(9):1228–1242. doi:10.1101/gr.252106.119

36. Cibulskis K, Lawrence MS, Carter SL, et al. Sensitive detection of somatic point mutations in impure and heterogeneous cancer samples. Nat Biotechnol. Mar 2013;31(3):213–9. doi:10.1038/nbt.2514

37. Lopez G, Egolf LE, Giorgi FM, Diskin SJ, Margolin AA. svpluscnv: analysis and visualization of complex structural variation data. Bioinformatics. Jul 27 2021;37(13):1912–1914. doi:10.1093/bioinformatics/btaa878

38. Oeck S, Malewicz NM, Hurst S, Al-Refae K, Krysztofiak A, Jendrossek V. The Focinator v2-0 - Graphical Interface, Four Channels, Colocalization Analysis and Cell Phase Identification. Radiat Res. Jul 2017;188(1):114–120. doi:10.1667/RR14746.1

39. Pinder J, Salsman J, Dellaire G. Nuclear domain ‘knock-in’ screen for the evaluation and identification of small molecule enhancers of CRISPR-based genome editing. Nucleic Acids Res. Oct 30 2015;43(19):9379–92. doi:10.1093/nar/gkv993

40. Bosse KR, Raman P, Zhu Z, et al. Identification of GPC2 as an Oncoprotein and Candidate Immunotherapeutic Target in High-Risk Neuroblastoma. Cancer Cell. Sep 11 2017;32(3):295–309 e12. doi:10.1016/j.ccell.2017.08.003

41. Laufer M, Nandula SV, Modi AP, et al. Structural requirements for the BARD1 tumor suppressor in chromosomal stability and homology-directed DNA repair. J Biol Chem. Nov 23 2007;282(47):34325–33. doi:10.1074/jbc.M705198200

42. Greenberg RA, Sobhian B, Pathania S, Cantor SB, Nakatani Y, Livingston DM. Multifactorial contributions to an acute DNA damage response by BRCA1/BARD1-containing complexes. Genes Dev. Jan 1 2006;20(1):34–46. doi:10.1101/gad.1381306

43. Rambhatla L, Chiu CP, Glickman RD, Rowe-Rendleman C. In vitro differentiation capacity of telomerase immortalized human RPE cells. Invest Ophthalmol Vis Sci. May 2002;43(5):1622–30.

44. Haeussler M, Schonig K, Eckert H, et al. Evaluation of off-target and on-target scoring algorithms and integration into the guide RNA selection tool CRISPOR. Genome Biol. Jul 5 2016;17(1):148. doi:10.1186/s13059-016-1012-2

45. Konishi H, Mohseni M, Tamaki A, et al. Mutation of a single allele of the cancer susceptibility gene BRCA1 leads to genomic instability in human breast epithelial cells. Proc Natl Acad Sci U S A. Oct 25 2011;108(43):17773–8. doi:10.1073/pnas.1110969108

46. Pathania S, Bade S, Le Guillou M, et al. BRCA1 haploinsufficiency for replication stress suppression in primary cells. Nat Commun. Nov 17 2014;5:5496. doi:10.1038/ncomms6496

47. Neu-Yilik G, Amthor B, Gehring NH, et al. Mechanism of escape from nonsense-mediated mRNA decay of human beta-globin transcripts with nonsense mutations in the first exon. RNA. May 2011;17(5):843–54. doi:10.1261/rna.2401811

48. Pathania S, Nguyen J, Hill SJ, et al. BRCA1 is required for postreplication repair after UV-induced DNA damage. Mol Cell. Oct 21 2011;44(2):235–51. doi:10.1016/j.molcel.2011.09.002

49. Farmer H, McCabe N, Lord CJ, et al. Targeting the DNA repair defect in BRCA mutant cells as a therapeutic strategy. Nature. Apr 14 2005;434(7035):917–21. doi:10.1038/nature03445

50. Takagi M, Yoshida M, Nemoto Y, et al. Loss of DNA Damage Response in Neuroblastoma and Utility of a PARP Inhibitor. J Natl Cancer Inst. Nov 1 2017;109(11) doi:10.1093/jnci/djx062

51. Raderschall E, Golub EI, Haaf T. Nuclear foci of mammalian recombination proteins are located at single-stranded DNA regions formed after DNA damage. Proc Natl Acad Sci U S A. Mar 2 1999;96(5):1921–6.

52. Newman EA, Chukkapalli S, Bashllari D, et al. Alternative NHEJ pathway proteins as components of MYCN oncogenic activity in human neural crest stem cell differentiation: implications for neuroblastoma initiation. Cell Death Dis. Dec 13 2017;8(12):3208. doi:10.1038/s41419-017-0004-9

53. Zhang W, Liu B, Wu W, et al. Targeting the MYCN-PARP-DNA Damage Response Pathway in Neuroendocrine Prostate Cancer. Clin Cancer Res. Feb 1 2018;24(3):696–707. doi:10.1158/1078-0432.CCR-17-1872

54. Ma X, Liu Y, Liu Y, et al. Pan-cancer genome and transcriptome analyses of 1,699 paediatric leukaemias and solid tumours. Nature. Mar 15 2018;555(7696):371–376. doi:10.1038/nature25795

55. Smith SA, Easton DF, Evans DG, Ponder BA. Allele losses in the region 17q12-21 in familial breast and ovarian cancer involve the wild-type chromosome. Nat Genet. Oct 1992;2(2):128–31. doi:10.1038/ng1092-128

56. Maxwell KN, Wubbenhorst B, Wenz BM, et al. BRCA locus-specific loss of heterozygosity in germline BRCA1 and BRCA2 carriers. Nat Commun. Aug 22 2017;8(1):319. doi:10.1038/s41467-017-00388-9

57. Wong M, Mayoh C, Lau LMS, et al. Whole genome, transcriptome and methylome profiling enhances actionable target discovery in high-risk pediatric cancer. Nat Med. Nov 2020;26(11):1742–1753. doi:10.1038/s41591-020-1072-4

58. McCarthy EE, Celebi JT, Baer R, Ludwig T. Loss of Bard1, the heterodimeric partner of the Brca1 tumor suppressor, results in early embryonic lethality and chromosomal instability. Mol Cell Biol. Jul 2003;23(14):5056–63. doi:10.1128/mcb.23.14.5056-5063.2003

59. Nutting C, Camplejohn RS, Gilchrist R, et al. A patient with 17 primary tumours and a germ line mutation in TP53: tumour induction by adjuvant therapy? Clin Oncol (R Coll Radiol). 2000;12(5):300–4.

60. Limacher JM, Frebourg T, Natarajan-Ame S, Bergerat JP. Two metachronous tumors in the radiotherapy fields of a patient with Li-Fraumeni syndrome. Int J Cancer. Aug 20 2001;96(4):238–42.

61. Ladenstein R, Potschger U, Pearson ADJ, et al. Busulfan and melphalan versus carboplatin, etoposide, and melphalan as high-dose chemotherapy for high-risk neuroblastoma (HR-NBL1/SIOPEN): an international, randomised, multi-arm, open-label, phase 3 trial. Lancet Oncol. Apr 2017;18(4):500–514. doi:10.1016/S1470-2045(17)30070-0

62. Desai AV, Heneghan MB, Li Y, et al. Toxicities of busulfan/melphalan versus carboplatin/etoposide/melphalan for high-dose chemotherapy with stem cell rescue for high-risk neuroblastoma. Bone Marrow Transplant. Sep 2016;51(9):1204–10. doi:10.1038/bmt.2016.84

63. Gillani R, Camp SY, Han S, et al. Germline predisposition to pediatric Ewing sarcoma is characterized by inherited pathogenic variants in DNA damage repair genes. Am J Hum Genet. Jun 2 2022;109(6):1026–1037. doi:10.1016/j.ajhg.2022.04.007

64. Kratz CP, Smirnov D, Autry R, et al. Heterozygous BRCA1/2 and Mismatch Repair Gene Pathogenic Variants in Children and Adolescents with Cancer. J Natl Cancer Inst. Aug 18 2022;doi:10.1093/jnci/djac151

