## Supplemental Methods and Data for "*BARD1* germline variants induce haploinsufficiency and DNA repair defects in neuroblastoma"

#### Cell culture

IMR-5 and RPE1 cells were obtained from the Children's Hospital of Philadelphia (CHOP) cell line bank. RPE1 cells are a human retinal pigment epithelium cell line immortalized through the retroviral insertion of human telomerase reverse transcriptase (hTERT) and were originally a kind gift from the laboratory of Dr. Michael Hogarty. Cell lines were cultured in RPMI containing 10% FBS and 2 mM L-Glutamine at 37°C under 5% $CO_2$. Cells were regularly tested for the presence of mycoplasma and genotyped to confirm cell identity using short tandem repeat (STR) typing.

#### Generation of isogenic cell models

IMR-5 and hTERT RPE1 cells were electroporated using a Lonza 4D-Nucleofector X-unit™ with 1.6 µg pU6-(BbsI)_CBh-Cas9-T2A-mCherry, into which one of four guide RNA sequences (R112*, R150*, E287fs, Q564*; **Supplemental Table 2**) had been cloned, and 0.4 µg single-stranded donor oligonucleotides containing the desired *BARD1* mutation and a synonymous PAM-ablating mutation. The pU6-(BbsI)_CBh-Cas9-T2A-mCherry plasmid was a gift from Ralf Kuehn (Addgene plasmid # 64324; http://n2t.net/addgene:64324; RRID:Addgene_64324).[1] Following electroporation, cells were transferred to media containing 5 µM L755507 (Selleck Chemicals) to enhance homology-directed repair efficiency.[2] Two days later, single mCherry-positive cells were sorted into 96-well plates using a BD FACSJazz cell sorter. Genomic DNA from single cell clones was extracted using the Qiagen DNeasy Blood and Tissue kit and the *BARD1* DNA was PCR amplified and Sanger sequenced to screen for the desired *BARD1* mutation. Heterozygous *BARD1* variants were confirmed using the PolyPeakParser program.[3] Clones that did not integrate a *BARD1* variant at either allele were also propagated for use as non-targeted control clones.

#### Quantitative RT-PCR

Total RNA was isolated from exponentially growing neuroblastoma cells utilizing RNeasy mini kits (Qiagen) and mRNAs were converted to cDNA using the SuperScript III system (ThermoFisher Scientific). Taqman® gene expression assays (Thermo Fischer Scientific) were used to quantitate *BARD1* (Hs00184427_m1 [*BARD1* exon 1-2 boundary] and Hs00957655_m1 [*BARD1* exon 9-10 boundary]), *BRCA1* (Hs00183233_m1), and *HPRT1* (Hs99999909_m1) on an Applied Biosystems 7900HT Sequence Detection System using standard cycling conditions. Relative transcript abundance was determined by the $2^{-\Delta\Delta Ct}$ method using *HPRT1* as an internal control.

**Immunofluorescence**

RPE1 *BARD1*$^{+/mut}$ and wild-type cells were seeded on poly-L-lysine coated coverslips (Electron Microscopy Sciences) and treated with 4 μM cisplatin or vehicle. Twenty-four hours after treatment, cells were fixed with 4% paraformaldehyde, stained with primary antibody (RAD51, Abcam ab88572, 1:100 or Phospho-Histone H2A.X (Ser139) (20E3), Cell Signaling Technology #9718, 1:800) followed by a secondary Alexa 488 or Alexa 555 antibody. Cells were mounted with ProLong gold with DAPI (Thermo Fisher Scientific, #P36931) and visualized with a Leica DM5000B microscope and photographed with a Leica DFC365 FX camera. RAD51 and γ-H2AX foci were quantified using Focinator v2.0 software and ImageJ.[4]

**mClover-LMNA assay**

IMR-5 *BARD1*$^{+/mut}$ and wild-type cells were co-transfected with 1.6 μg pX330-LMNA-gRNA1 and 0.4 μg pCR2.1 Clover-LMNA using the Lonza 4D-Nucleofector X-unit™ system. The mClover-LMNA reagents were a kind gift form the laboratory of Graham Dellaire.  After 3 days, cells were fixed in 2% paraformaldehyde and analyzed on a CytoFLEX-LX flow cytometer to quantitate clover-positive cells.

**Cytotoxicity studies**

IMR-5 and RPE1 *BARD1*$^{+/mut}$ cells and paired wild-type cells were plated on Day 1 in a 96-well plate. On Day 2, serial dilutions of olaparib (Selleck Chemicals, DMSO) or cisplatin (Selleck Chemicals, $H_2O$) were added. After 4 days, cell viability was determined using a CellTiter-Glo® Assay (Promega) in a GloMax (Promega) plate reader according to the manufacturer's instructions. Luminescence values were normalized to vehicle treated wells and data were analyzed and graphed in GraphPad Prism software and a log (inhibitor) vs. response nonlinear regression model was used to calculate $IC_{50}$s.

### *In vivo* IMR5 *BARD1*$^{+/mut}$ xenograft efficacy studies

*In vivo* murine xenograft efficacy studies were designed to assess the efficacy of olaparib in IMR5 *BARD1*$^{+/mut}$ isogenic cell line derived xenograft models. IMR5 *BARD1*$^{+/mut}$ isogenic cell lines were expanded *in vitro* and 5 x $10^6$ cells were mixed with Matrigel (Corning, cat# 354234) and injected into the flanks of CB17-SCID mice (Taconic Farms, Germantown NY). When the tumors reached a size of 1-1.5 $cm^3$, they were serially passaged into study CB17-SCID mice. When tumors reached enrollment size (0.15 $cm^3$-0.3 $cm^3$), mice were then randomly enrolled into 2 treatment cohorts (Olaparib or vehicle; n=10 per cohort), using a rolling enrollment to ensure almost identical tumor sizes across treatment cohorts. Olaparib was dosed intraperitoneally at 20mg/kg once daily for 28 days. Tumor sizes were measured at least twice weekly using calipers and tumor volumes were calculated as: volume = ((diameter1/2 + diameter2/2)3*0.5236)/1000. Mice weights were also measured at least twice weekly, and mice were monitored daily for signs of any clinical toxicity. Mice were sacrificed when tumor burden reached 2 $cm^3$ or they showed any signs of distress including excessive weight loss. All *in vivo* animal studies were performed according to Children's Hospital of Philadelphia (CHOP) policies in the Department of Veterinary Research (DVR) and were conducted according to an approved IACUC Protocol (#0006430). Up to 5 mice were maintained in cages under barrier conditions in a pathogen-free facility fully accredited by the Association for Assessment and Accreditation of Laboratory Animal Care (AAALAC).

### Quantification and statistical analysis

Differences between groups were presented as the mean ± SEM as noted in the figure legends. Experimental sample numbers (n) are indicated in the figures, figure legends and results section. All statistical analysis was done with GraphPad Prism and p-values < 0.05 were considered statistically significant. RAD51 and γ-H2AX foci were quantified using Focinator v2.0 software and ImageJ.[4]

### Whole-genome sequencing of IMR-5 cells

All code is available on GitHub (https://github.com/diskin-lab-chop/nbl-bard1), except when a public pipeline is referenced. After 20 passages, genomic DNA was extracted from three IMR-5 *BARD1*$^{+/mut}$ clonal cell lines and one non-targeted control clone using the Qiagen Blood and Tissue kit and then treated with RNase to digest RNA. DNA integrity was assessed by pulse-field gel electrophoresis. Libraries were prepared with a 1%

PhiX spike-in, fragmented, and sequenced on an Illumina HiSeq 10X using S2 chemistry with 150 bp paired-end reads to at least 30X mean coverage. Separately, DNA from WT parental IMR-5 cells (prior to 20 passages) was isolated and sequenced with similar methods, and this parental sample served as the "normal" control for filtering variant calls from the *BARD1*$^{+/mut}$ and non-targeted control clones. FASTQ files were aligned against hg19 (b37 reference from the Broad Institute) with BWA-MEM 0.7.17[5] using the public Seven Bridges Genomics workflow "Whole Genome Sequencing - BWA + GATK 4.0 (with Metrics)" on CAVATICA (https://www.cavatica.org/, app ID: admin/sbg-public-data/whole-genome-sequencing-bwa-gatk-4-0, revision 41). After alignment, BAM files were randomly downsampled with Picard DownsampleSam to achieve 50x mean coverage, or 1.1 billion aligned reads, for each of the three *BARD1*$^{+/mut}$ clones and the non-targeted control clone. Only chromosomes 1-22, X, and Y were considered for subsequent analyses.

**Copy number analysis**

Copy number segmentation profiles were generated with Control-FREEC v11.5[6] using a public Seven Bridges workflow on CAVATICA (app ID: admin/sbg-public-data/control-freec-11-5, revision 4) with default settings. The parental IMR-5 cell line (described above) was used as the normal control for paired analysis. Segments containing less than 5 genomic bins (approximately 5.6 kb) were removed. Segments overlapping 50% or more with the ENCODE hg19 blacklist[7] or segmental duplications (as defined by the UCSC Genome Browser[8], considering only those with >95% identity) were removed. Copy number ratio thresholds for gain and loss were set at 1.2 and 0.8, respectively. Breakpoint analysis was performed with the *svpluscnv* R package (https://github.com/ccbiolab/svpluscnv)[9], based on methods developed by Lopez *et al*.[10] Double-strand breaks were quantified by counting regions where the fold change between any two adjacent segments was greater than 1.2 or less than 0.8 (*fc.pct*=0.2).

**Structural variant (SV) analysis**

SVs were called with Delly v0.7.9[11] in paired mode, using the parental IMR-5 cell line as the normal control. SVs were filtered for the default PASS criteria at the dataset and individual levels and required to have at least 5 reads supporting the alternate allele (considering both split-read and paired-read support). SVs with one or more breakpoints falling within the ENCODE blacklist or segmental duplications (described above) were removed. Stringent filtering (shown in **Figure 3B-D, Supplemental Figure 2B-C**) considered only precise SVs supported by split reads, whereas relaxed filtering (shown in **Supplemental Figure 3A-E**) included both precise and imprecise SVs.

**Single-nucleotide variant (SNV) and indel analysis**

SNVs and indels were called with MuTect2[12] from GATK v4.1.3.0, again using parental IMR-5 as the normal control. The read orientation bias filter was applied. Variants flagged by FilterMutectCalls for any reason except "clustered_events" were removed. Di- and tri-nucleotide polymorphism calls were removed. For all figures except the mutational signature analysis, variants were required to have at least 5 reads supporting the alternate allele.

**Mutational signature analysis**

The above filtered variant calls were used as input to the deconstructSigs v. 1.9.0 R package to perform mutational signature analysis using the following signature sets: COSMIC v2 SBS and COSMIC v3.2 SBS (https://cancer.sanger.ac.uk/signatures/). The v3.2 COSMIC mutational signatures were down-sampled to remove signatures driven by therapy, environmental exposures, and/or sequencing artifacts, along with SBS39 due to the high similarity to SBS3, while maintaining other neuroblastoma-specific[13] and biologically relevant signatures. Our analysis code can be found on GitHub (https://github.com/diskin-lab-chop/nbl-bard1).

### Supplemental Data

**Supplemental Table 1.** Characteristics of neuroblastoma-associated germline *BARD1* variants.

| USI | Age at diagnosis (Days) | Sex | *MYCN* | Risk group | Variant | Exon | Cell line models | Other cancers associated with germline *BARD1* variant |
|---|---|---|---|---|---|---|---|---|
| PATZRU | 833 | Male | NA | High | c.159-1G>T (splice site) | 2 | | Breast[14] |
| PAHYWC | 704 | Male | Amp | High | c.C334T; p.R112* | 3 | IMR-5 x 2 | Breast[14, 15] |
| PARSEA | 1779 | Male | NA | High | c.448C>T; p.R150* | 4 | IMR-5 | Breast[14, 16]<br>Ovarian[17] |
| PATHJZ | 340 | Female | NA | Intermediate | c.860_861del; p.E287fs | 4 | IMR-5 x 3 | - |
| PASGEE | 1825 | Male | NA | High | c.1677+1G>T (splice donor) | 7 | | - |
| PASFDU | 758 | Female | NA | High | c.C1690T; p. Q564* | 8 | RPE1 | Breast[14, 16, 18-22]<br>Ovarian[16, 19, 23, 24]<br>Endometrial[25]<br>Colorectal[22, 26] |
| PATGWT | 591 | Male | Amp | High | c.1921C>T; p.R641* | 10 | | Breast[21, 27, 28]<br>Pancreatic[29] |
| PASCIX | 1660 | Male | NA | High | c.1935_1954dup; p.Glu652fs | 10 | | Breast[16, 30, 31] |

Amp., *MYCN* amplified tumor; NA, *MYCN* non-amplified tumor.

**Supplemental Table 2.** Guide RNAs and repair template oligonucleotides used to generate *BARD1* isogenic. cell lines.

| Variant | Guide RNA | Single-stranded repair oligonucleotide | Notes |
|---|---|---|---|
| c.C334T; p.R112* | CTTGAAGATAAATAGACAAC<br>GTTGTCTATTTATCTTCAAG | ACTGATGAATTTAACTAAGAGAGATAGGGATAGTT<br>CTTACCTGACAGCTCATTG<br>TCATGTAGCAAATTTCAAAGCTTACTACAAAGTTGA<br>ATCATGCTGTC*G*AGTTGTC<br>TATTTATCTTCAAGTCTTGTATCCAGGCCGGG | |
| c.C448T; p.R150* | ATCTGACTTTCTTACTTCGA<br>TCGAAGTAAGAAAGTCAGAT | GCATCTTTTTTTATTGCAGGCTGGGTTTGCACTGA<br>AGCTTTACTCACAACATAT<br>CTGACTTTCTTACTTCAAGG*AGA*AAACCACATTTTA<br>ATTGAATTCTTCTTGTTTC<br>CTGCATCATTAAACAAACTTTTCCTAGGTTTA | |
| c.860_861del; p.E287fs | AGTCTCCAGACACTAAGAGC<br>GCTCTTAGTGTCTGGAGACT | GGCTCCTTGACAGAATCTGAATGTTTTGGAAGTTT<br>AACTGAAGTCTCTTTACCA<br>TTGGCTGAGCAAATAG__TCTCCAGACACTAAGAG<br>CAG*A*AATGAAGTAGTGACT<br>CCTGAGAAGGTCTGCAAAAATTATCTTACATC | Utilzed for E287 #1 |
| | | TAGATGTAAGATAATTTTTGCAGACCTTCTCAGGA<br>GTCACTACTTCATTC*T*[a]TGC<br>TCTTAGTGTCTGGAGA__CTATTTGCTCAGCCAAT<br>GGTAAAGAGACTTCAGTT<br>AAACTTCCAAAACATTCAGATTCTGTCAAGGAGCC | Utilized for E287 #2,3 |
| c.C1960T; p.Q564* | TATATTAACAGATGAACACT<br>AGTGTTCATCTGTTAATATA | TCACTGAGCATTTTCTGTTGTTCTGAAGACAGCCC<br>ACTGCCTATAAGTACAAGA<br>GGTCCATCCCTACGCTA*T*CCAGTGTTCATCTGTTA<br>ATATAAAAGGAGATACCAGTGTTAAAAACATTAGA<br>CGACTAGACAAAGACAT | |

[a]This PAM variant is non-synonymous, but occurs after the frameshift at codon 287 and subsequent truncating variant at codon 291
Double underline, Pathogenic variant; *Italic*: Protospacer adjacent motif (PAM) variant.

**Supplemental Table 3.** Possible CRISPR off-target sites evaluated via Sanger sequencing.

| Guide RNA | Type | Forward Primer | Reverse Primer | CFD Score |
|---|---|---|---|---|
| R112* | Intergenic | ACCTCACATGTGCTAAGGATGT | GTGATTTTCCTTACGAAGTGCTGA | 0.90 |
| | Exon (RP6) | AGGTCTTACTCCCAAAACATGTCA | ACATGCAAAGTAAACACTTGCA | 0.13 |
| | Exon (RP11) | AGCTTTTACACATGCTGAGACT | CACACACACAAACACCACACA | 0.07 |
| R150* | Intergenic | AGGGCAAGACAAGACTGCAA | CTTGGCTGGAAGGAGCATGA | 0.41 |
| | Exon (EPAS1) | TGGTTCTCTGGCCATTTCCC | CAAATGTGAGGTGCTGCCAC | 0.14 |
| E287fs | Intergenic | GCATTTTAGCATGGTGTCTATGGT | ACGTATCAACAAATAGCATTCACT | 0.67 |
| | Exon (CCR9) | TGTTATCGGGTAGCTGCCTG | GATGCAACTCTCCCTGGGAC | 0.41 |
| | Exon (LL22NC03) | TCCTGTCGTGTCTGTTTCGG | GAGCCACAGGTGAGAGTGAC | 0.05 |
| Q564* | Intergenic | TCATTGAACTGCATACAAGTGCT | ATTGAAAACTGGATATTCTCTGCTT | 0.36 |
| | Exon (RP11) | CCTGGGACTCGAACCGTATG | GTACAACCTGGTGTGGAGGG | 0.33 |
| | Exon (UBE2G1) | AAAGCCACCTCGTTCAGTGT | ACTTCCCTTCCTCTGTCGGA | 0.04 |

### Supplemental Figures and Figure Legends

#### Supplemental Figure 1

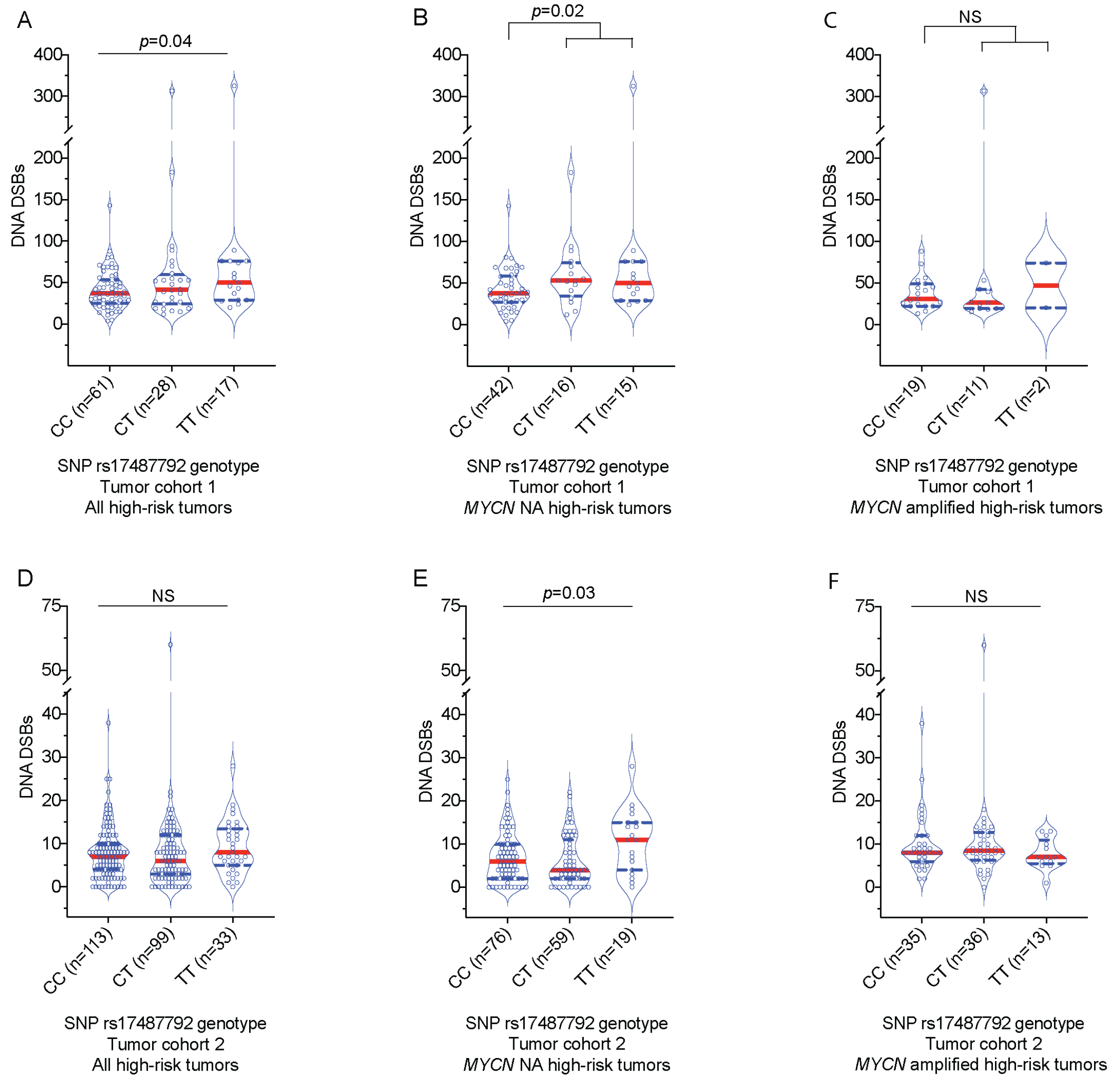


**Supplemental Figure 1. Common *BARD1* germline risk variants correlate with genome-wide deficiencies in DNA repair in high-risk *MYCN* non-amplified primary neuroblastomas.**

(**A**-**F**) Violin plots depicting the number of DNA DSBs in neuroblastoma tumors from only high-risk patients with different germline SNP rs174877792 genotypes. Panels **A**, **B** and **C** depict DNA DSBs in all high-risk tumors, high-risk tumors without *MYCN* amplification and high-risk tumors with *MYCN* amplification in tumor cohort 1, respectively. Panels **D**, **E** and **F** depict DNA DSBs in all high-risk tumors, high-risk tumors without *MYCN* amplification and high-risk tumors with *MYCN* amplification in tumor cohort 2, respectively. Red dotted line denotes median and blue dotted lines denotes quartiles.

*MYCN* NA, *MYCN* non-amplified.

Associated with **Figure 1**.

**Supplemental Figure 2**

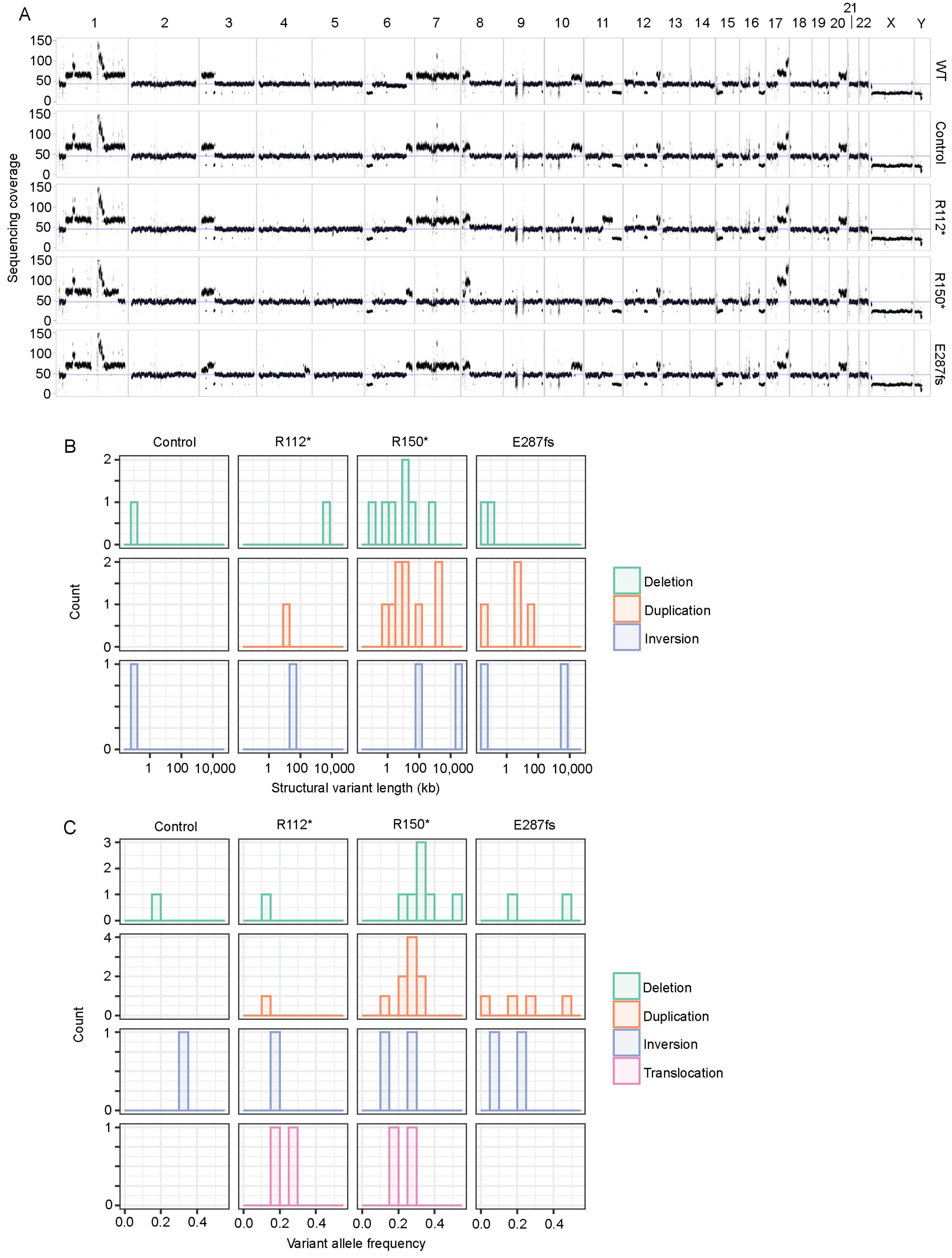

**Supplemental Figure 2.** ***BARD1***$^{+/mut}$ **neuroblastoma IMR-5 clonal cell lines exhibit genome-wide genomic instability.**

(**A**) Whole-genome sequencing coverage for WT parental cells, a non-targeted control clone and *BARD1*$^{+/mut}$ isogenic IMR-5 cells.

(**B**) Histograms showing length of structural variants in control and *BARD1*$^{+/mut}$ isogenic IMR-5 cells.

(**C**) Histograms showing allele frequency of structural variants in control and *BARD1*$^{+/mut}$ isogenic IMR-5 cells.

Associated with **Figure 3**.

### Supplemental Figure 3

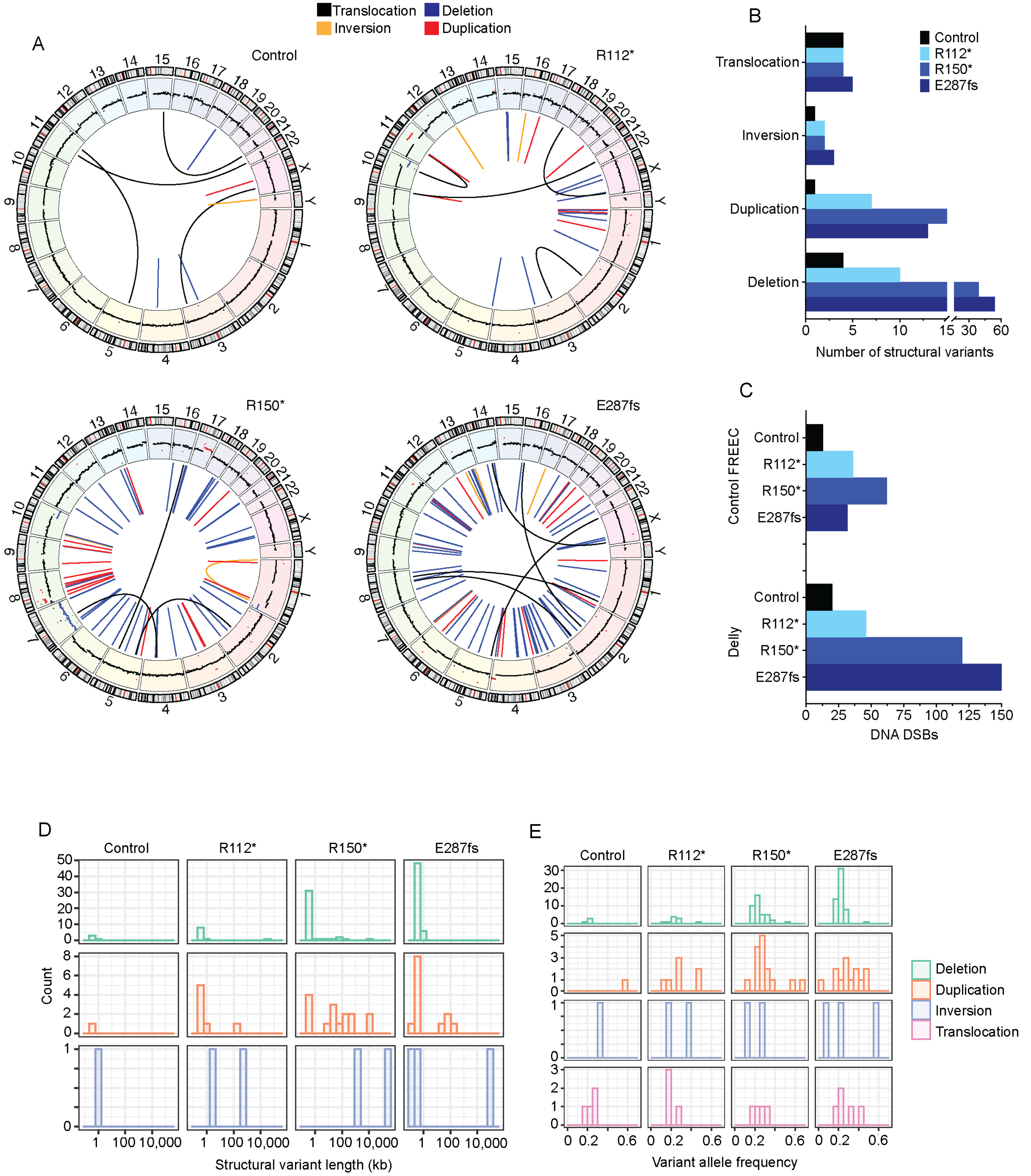

**Supplemental Figure 3. Structural variant analysis with relaxed filtering confirms increased genome instability in *BARD1*$^{+/mut}$ neuroblastoma IMR-5 clonal cell lines.**

(**A**) Circos plots depicting identified structural variants in control and *BARD1*$^{+/mut}$ isogenic IMR-5 models using less stringent filtering parameters.

(**B**) Counts of structural variants in control and *BARD1*$^{+/mut}$ isogenic IMR-5 cells using less stringent filtering parameters.

(**C**) Counts of DNA DSBs in control and BARD1$^{+/mut}$ IMR-5 cells, quantified from the Control-FREEC copy number (**top**) and the Delly structural variant data (**bottom**) using less stringent filtering parameters.

(**D**) Histograms showing length of structural variants in control and *BARD1*$^{+/mut}$ isogenic IMR-5 cells using less stringent filtering parameters.

(**E**) Histograms showing allele frequency of structural variants in control and *BARD1*$^{+/mut}$ IMR-5 isogenic cells using less stringent filtering parameters.

Associated with **Figure 3**.

### Supplemental Figure 4

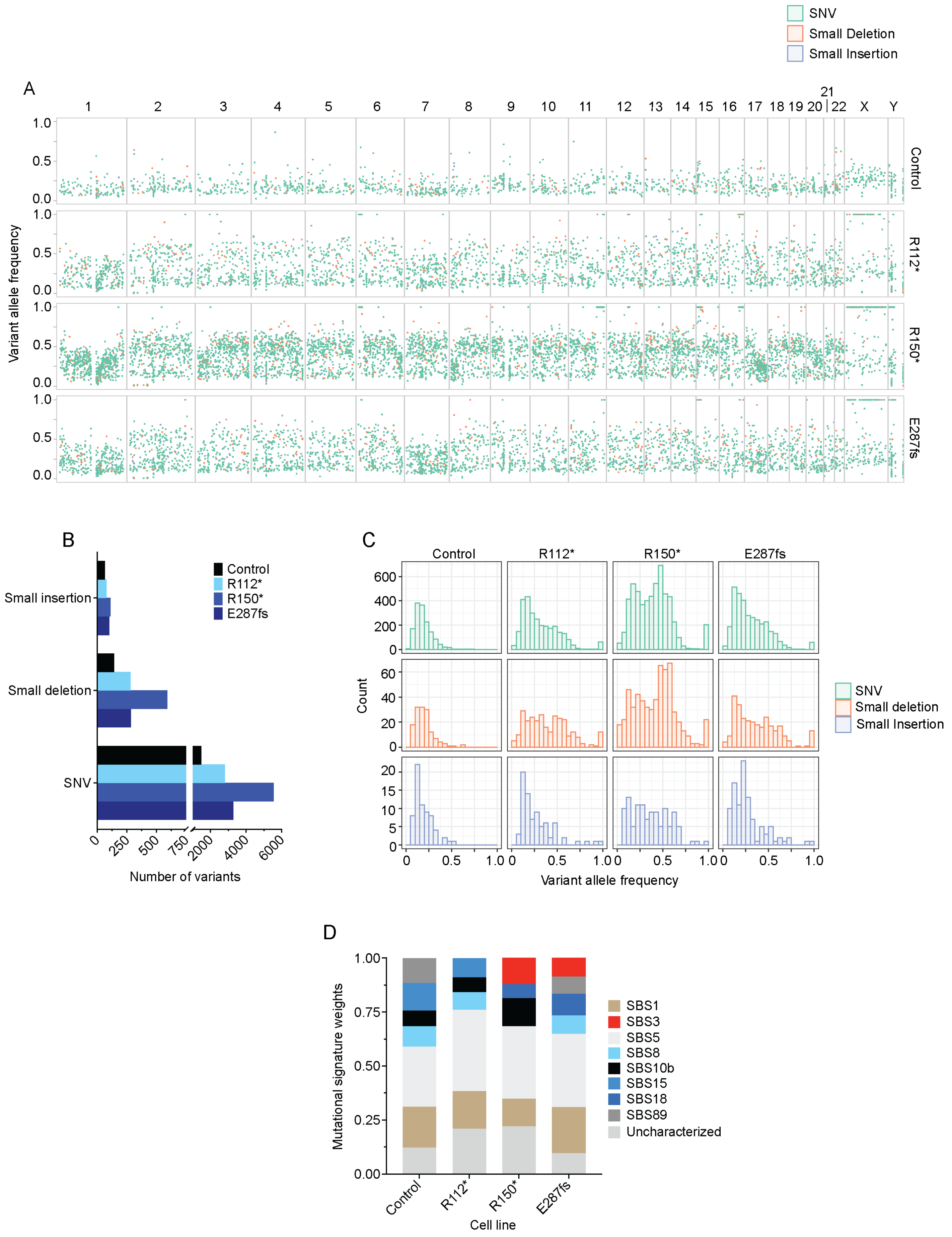

**Supplemental Figure 4. IMR-5 *BARD1*$^{+/mut}$ isogenic cells acquired more SNVs and indels than the non-targeted control clone.**

(**A**) Variant allele frequency distribution across the genome for SNVs and indels acquired in control clone and *BARD1*$^{+/mut}$ isogenic IMR-5 cells relative to WT parental IMR-5 cells.

(**B**) Count of SNVs and indels identified in control and *BARD1*$^{+/mut}$ isogenic IMR-5 cells.

(**C**) Histograms showing allele frequency of SNVs and indels identified in control and *BARD1*$^{+/mut}$ isogenic IMR-5 models.

(**D**) Plot of mutational signature weights in non-targeted control and *BARD1*$^{+/mut}$ IMR-5 cells using COSMIC mutational signatures (v3.2).

Associated with **Figure 3**.

**Supplemental Figure 5**

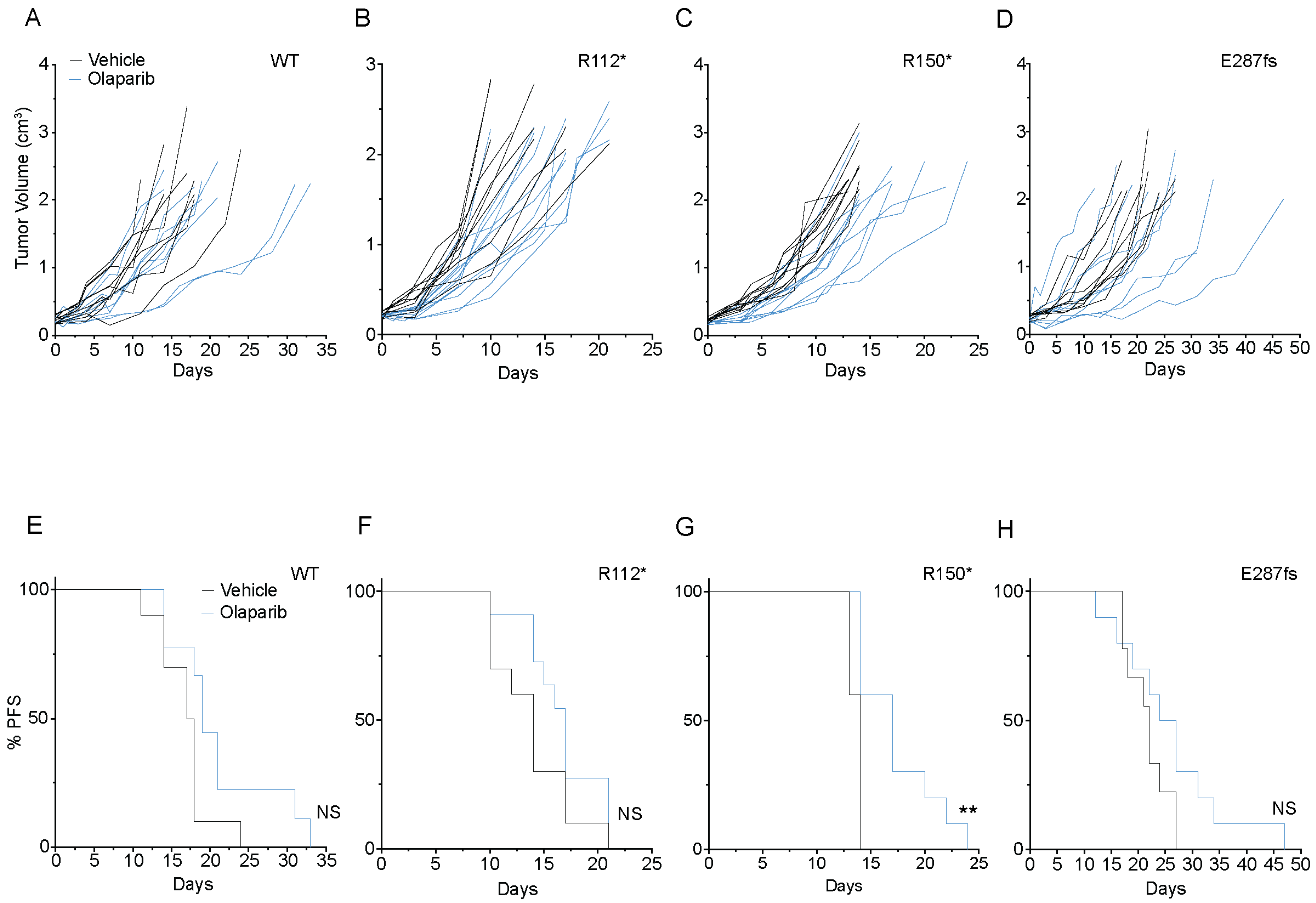


**Supplemental Figure 5. IMR-5 and RPE1 *BARD1*$^{+/mut}$ models show increased sensitivity to olaparib and cisplatin.**

(**A**-**D**) Individual tumor growth curves of WT and *BARD1*$^{+/mut}$ IMR-5 xenografts treated with daily olaparib or vehicle [WT IMR-5 (**A**), *BARD1*$^{+/R112*}$ (**B**), *BARD1*$^{+/R150*}$ (**C**), *BARD1*$^{+/E287fs}$ (**D**)].

(**E**-**H**) Progression-free survival of mice with WT and *BARD1*$^{+/mut}$ IMR-5 xenografts treated with daily olaparib or vehicle [WT IMR-5 (**E**), *BARD1*$^{+/R112*}$ (**F**), *BARD1*$^{+/R150*}$ (**G**), *BARD1*$^{+/E287fs}$ (**H**)].

***P* < 0.01; NS, not significant.

Associated with **Figure 4**.

**Supplemental References**


1. Chu VT, Weber T, Wefers B, et al. Increasing the efficiency of homology-directed repair for CRISPR-Cas9-induced precise gene editing in mammalian cells. *Nat Biotechnol*. May 2015;33(5):543-8. doi:10.1038/nbt.3198

2. Yu C, Liu Y, Ma T, et al. Small molecules enhance CRISPR genome editing in pluripotent stem cells. *Cell Stem Cell*. Feb 5 2015;16(2):142-7. doi:10.1016/j.stem.2015.01.003

3. Hill JT, Demarest BL, Bisgrove BW, Su YC, Smith M, Yost HJ. Poly peak parser: Method and software for identification of unknown indels using sanger sequencing of polymerase chain reaction products. *Dev Dyn*. Dec 2014;243(12):1632-6. doi:10.1002/dvdy.24183

4. Oeck S, Malewicz NM, Hurst S, Al-Refae K, Krysztofiak A, Jendrossek V. The Focinator v2-0 - Graphical Interface, Four Channels, Colocalization Analysis and Cell Phase Identification. *Radiat Res*. Jul 2017;188(1):114-120. doi:10.1667/RR14746.1

5. Li H, Durbin R. Fast and accurate short read alignment with Burrows-Wheeler transform. *Bioinformatics*. Jul 15 2009;25(14):1754-60. doi:10.1093/bioinformatics/btp324

6. Boeva V, Popova T, Bleakley K, et al. Control-FREEC: a tool for assessing copy number and allelic content using next-generation sequencing data. *Bioinformatics*. Feb 1 2012;28(3):423-5. doi:10.1093/bioinformatics/btr670

7. Amemiya HM, Kundaje A, Boyle AP. The ENCODE Blacklist: Identification of Problematic Regions of the Genome. *Sci Rep*. Jun 27 2019;9(1):9354. doi:10.1038/s41598-019-45839-z

8. Karolchik D, Hinrichs AS, Furey TS, et al. The UCSC Table Browser data retrieval tool. *Nucleic Acids Res*. Jan 1 2004;32(Database issue):D493-6. doi:10.1093/nar/gkh103

9. Lopez G, Egolf LE, Giorgi FM, Diskin SJ, Margolin AA. svpluscnv: analysis and visualization of complex structural variation data. *Bioinformatics*. Jul 27 2021;37(13):1912-1914. doi:10.1093/bioinformatics/btaa878

10. Lopez G, Conkrite KL, Doepner M, et al. Somatic structural variation targets neurodevelopmental genes and identifies SHANK2 as a tumor suppressor in neuroblastoma. *Genome Res*. Sep 2020;30(9):1228-1242. doi:10.1101/gr.252106.119

11. Rausch T, Zichner T, Schlattl A, Stutz AM, Benes V, Korbel JO. DELLY: structural variant discovery by integrated paired-end and split-read analysis. *Bioinformatics*. Sep 15 2012;28(18):i333-i339. doi:10.1093/bioinformatics/bts378

12. Cibulskis K, Lawrence MS, Carter SL, et al. Sensitive detection of somatic point mutations in impure and heterogeneous cancer samples. *Nat Biotechnol*. Mar 2013;31(3):213-9. doi:10.1038/nbt.2514

13. Brady SW, Liu Y, Ma X, et al. Pan-neuroblastoma analysis reveals age- and signature-associated driver alterations. *Nat Commun*. Oct 14 2020;11(1):5183. doi:10.1038/s41467-020-18987-4

14. Weber-Lassalle N, Borde J, Weber-Lassalle K, et al. Germline loss-of-function variants in the BARD1 gene are associated with early-onset familial breast cancer but not ovarian cancer. *Breast Cancer Res*. Apr 29 2019;21(1):55. doi:10.1186/s13058-019-1137-9

15. Gonzalez-Rivera M, Lobo M, Lopez-Tarruella S, et al. Frequency of germline DNA genetic findings in an unselected prospective cohort of triple-negative breast cancer patients participating in a platinum-based neoadjuvant chemotherapy trial. *Breast Cancer Res Treat*. Apr 2016;156(3):507-515. doi:10.1007/s10549-016-3792-1

16. Susswein LR, Marshall ML, Nusbaum R, et al. Pathogenic and likely pathogenic variant prevalence among the first 10,000 patients referred for next-generation cancer panel testing. *Genet Med*. Aug 2016;18(8):823-32. doi:10.1038/gim.2015.166

17. Norquist BM, Harrell MI, Brady MF, et al. Inherited Mutations in Women With Ovarian Carcinoma. *JAMA Oncol*. Apr 2016;2(4):482-90. doi:10.1001/jamaoncol.2015.5495

18. Domagala P, Jakubowska A, Jaworska-Bieniek K, et al. Prevalence of Germline Mutations in Genes Engaged in DNA Damage Repair by Homologous Recombination in Patients with Triple-Negative and Hereditary Non-Triple-Negative Breast Cancers. *PLoS One*. 2015;10(6):e0130393. doi:10.1371/journal.pone.0130393

19. Klonowska K, Ratajska M, Czubak K, et al. Analysis of large mutations in BARD1 in patients with breast and/or ovarian cancer: the Polish population as an example. *Sci Rep*. May 21 2015;5:10424. doi:10.1038/srep10424

20. Ratajska M, Antoszewska E, Piskorz A, et al. Cancer predisposing BARD1 mutations in breast-ovarian cancer families. *Breast Cancer Res Treat*. Jan 2012;131(1):89-97. doi:10.1007/s10549-011-1403-8

21. De Brakeleer S, De Greve J, Desmedt C, et al. Frequent incidence of BARD1-truncating mutations in germline DNA from triple-negative breast cancer patients. *Clin Genet*. Mar 2016;89(3):336-40. doi:10.1111/cge.12620

22. Adamovich AI, Banerjee T, Wingo M, et al. Functional analysis of BARD1 missense variants in homology-directed repair and damage sensitivity. *PLoS Genet*. Mar 2019;15(3):e1008049. doi:10.1371/journal.pgen.1008049

23. Ratajska M, Matusiak M, Kuzniacka A, et al. Cancer predisposing BARD1 mutations affect exon skipping and are associated with overexpression of specific BARD1 isoforms. *Oncol Rep*. Nov 2015;34(5):2609-17. doi:10.3892/or.2015.4235

24. Ramus SJ, Song H, Dicks E, et al. Germline Mutations in the BRIP1, BARD1, PALB2, and NBN Genes in Women With Ovarian Cancer. *J Natl Cancer Inst*. Nov 2015;107(11)doi:10.1093/jnci/djv214

25. Ring KL, Bruegl AS, Allen BA, et al. Germline multi-gene hereditary cancer panel testing in an unselected endometrial cancer cohort. *Mod Pathol*. Nov 2016;29(11):1381-1389. doi:10.1038/modpathol.2016.135

26. Blazer KR, Nehoray B, Solomon I, et al. Next-Generation Testing for Cancer Risk: Perceptions, Experiences, and Needs Among Early Adopters in Community Healthcare Settings. *Genet Test Mol Biomarkers*. Dec 2015;19(12):657-65. doi:10.1089/gtmb.2015.0061

27. Gass J, Tatro M, Blackburn P, Hines S, Atwal PS. BARD1 nonsense variant c.1921C>T in a patient with recurrent breast cancer. *Clin Case Rep*. Feb 2017;5(2):104-107. doi:10.1002/ccr3.793

28. Feliubadalo L, Tonda R, Gausachs M, et al. Benchmarking of Whole Exome Sequencing and Ad Hoc Designed Panels for Genetic Testing of Hereditary Cancer. *Sci Rep*. Jan 4 2017;7:37984. doi:10.1038/srep37984

29. Hu C, Hart SN, Bamlet WR, et al. Prevalence of Pathogenic Mutations in Cancer Predisposition Genes among Pancreatic Cancer Patients. *Cancer Epidemiol Biomarkers Prev*. Jan 2016;25(1):207-11. doi:10.1158/1055-9965.EPI-15-0455

30. Couch FJ, Hart SN, Sharma P, et al. Inherited mutations in 17 breast cancer susceptibility genes among a large triple-negative breast cancer cohort unselected for family history of breast cancer. *J Clin Oncol*. Feb 1 2015;33(4):304-11. doi:10.1200/JCO.2014.57.1414

31. De Brakeleer S, De Greve J, Loris R, et al. Cancer predisposing missense and protein truncating BARD1 mutations in non-BRCA1 or BRCA2 breast cancer families. *Hum Mutat*. Mar 2010;31(3):E1175-85. doi:10.1002/humu.21200
